# A general approach to reduce off-target radioactivity in vivo via Tetrazine-Knock-Out (TKO)

**DOI:** 10.1101/2024.05.30.596510

**Authors:** Swarbhanu Sarkar, Jonathan M. Pham, Kimberly J. Edwards, Nitika Sharma, Kexiang Xu, A. Paden King, Andres Fernandez del Castillo, Michael D. Farwell, Daniel A. Pryma, Stephen J. Schuster, Mark A. Sellmyer

## Abstract

Monoclonal antibodies have had a remarkable impact on cancer therapy due to their high target specificity. However, their large molecular weight results in slow blood clearance, which can take weeks to clear from circulation. As companion nuclear imaging and diagnostic tools, these characteristics force delayed imaging and the use of isotopes with long half-lives such as ^89^Zr. For optimal clinical application, it is desirable that radioimmunoconjugates remain in the blood for just enough time to accumulate adequately in target tissues, while non-targeted or circulating radioactivity is ideally rapidly excreted from the body to maximize imaging contrast and minimize radiation dose to healthy tissues. We addressed the current challenges of antibody-based imaging by developing rituximab radioimmunoconjugates that accumulate sufficient activity for tumor imaging within 24 h of administration, while clearing circulating radioactivity via administration of a small molecule clearing agent. Rituximab, an anti-CD20 monoclonal antibody, is used as standard first-line therapy for diffuse large B-cell lymphoma. CD20 is expressed by 95% of B-lymphocytes and their malignant counterparts, making it a therapeutic target for B-cell malignancies. We attached ^125^I, ^68^Ga, and ^89^Zr to rituximab using a “clickable” linker containing *trans*-cyclooctene and tested the ability of tetrazines to induce the inverse electron demand Diels-Alder reaction (iEDDA) after antibody administration. This “tetrazine-knock-out” (TKO) approach liberates the radioactivity from rituximab in the bloodstream, resulting in its rapid renal excretion which enhances target-to-background ratios, and minimizes off-target radiation exposure. Due to the internalization of the radioimmunoconjugate in CD20^+^ tumor cells, no substantial clearance was observed from Raji xenografts. We characterized different leaving groups, several cellular models and antibodies with distinct internalizaing properties. The TKO approach opens opportunities to use radiolabeled antibodies for low-abundance or heterogeneously expressed biologic targets and may allow radioimmunotherapy (RIT) for targets traditionally untenable due to dose-limiting toxicities.

**Graphical abstract:** 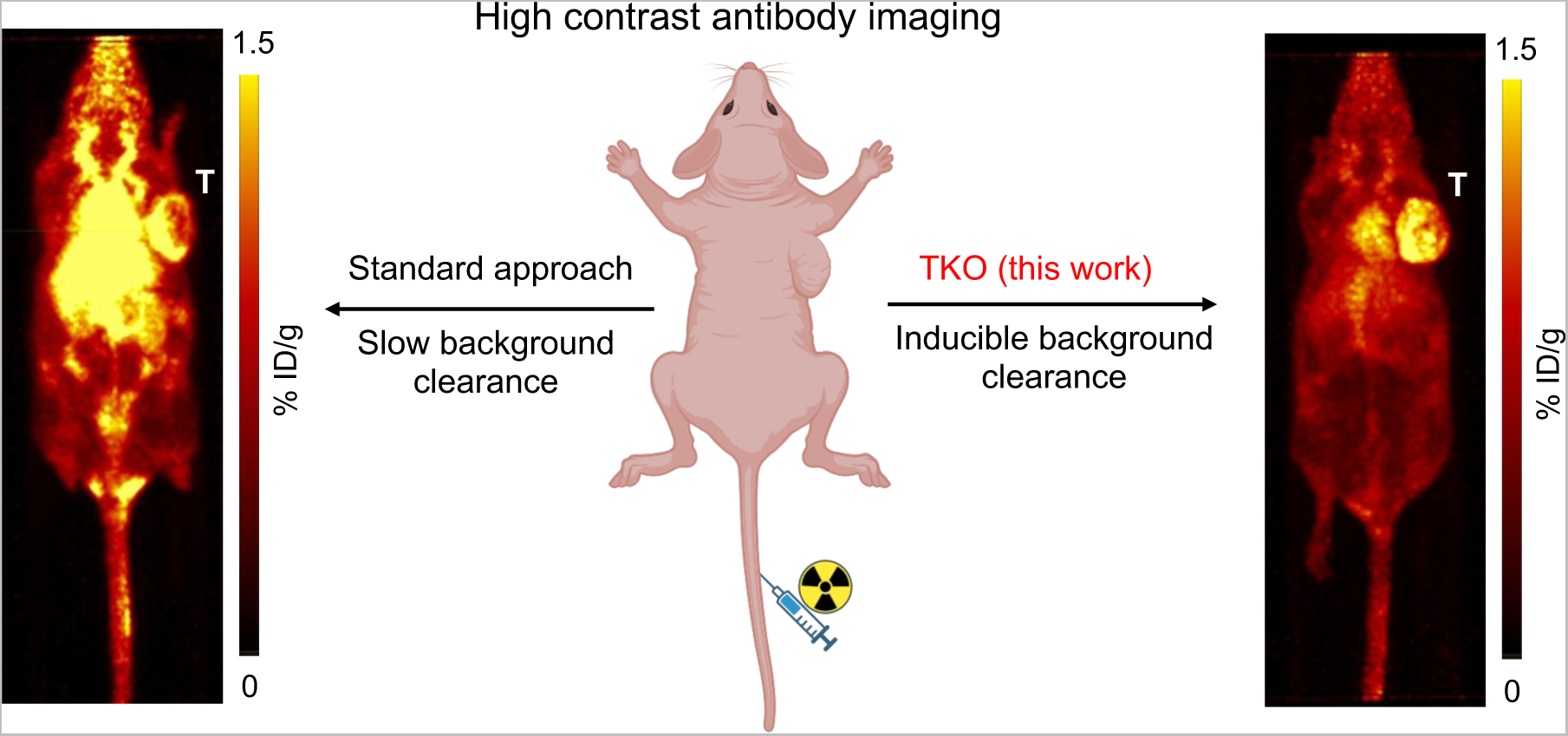

## Introduction

Monoclonal antibodies (mAbs) have had a remarkable impact on cancer therapy.^1–3^ Currently, over 570 antibody-based therapeutics are in clinical trials, including 62 in late-stage clinical studies, and 79 therapeutic mAbs have been approved by the United States Food and Drug Administration.^4^ Antibodies such as atezoliumab (an anti-PD-L1 therapy) have also been labeled with radioisotopes including ^89^Zr in early phase clinical trials, demonstrating the potential of such radiopharmaceuticals to guide therapy choice and predict therapeutic response.^5^ While offering the promise of quantitative measurements of antigen density and greater specificity for tumor cells,^6^ a substantial drawback emerges in the extended timeframe required to achieve optimal target-to-background signals.^7–8^ When used as companion imaging and diagnostic tools, the high molecular weight of antibodies results in slow blood clearance, which can take weeks to clear from circulation. In both preclinical and clinical settings, the optimal imaging contrast typically becomes apparent 3-7 days after injection. Although tumor accumulation can rapidly reach substantial levels (approximately 10% of injected activity per gram of tumor in mice) within a few hours post-injection, high levels of circulating radioactivity persist for days.^9–10^ This prolonged circulation not only adds to background and blood pool signal, but also increases the radiation dose to healthy organs which limits the amount of radioactivity that can be injected – two critical components ultimately resulting in low-quality images.^11–12^ Thus, there is a gap in our ability to image radio-immunoconjugates on the same or very next day after administration, in contrast to small molecule radiotracers like [^18^F]FDG or ligands targeting the prostate specific membrane antigen (PSMA) and the somatostatin receptor.^13^ The extended delay in imaging time also restricts immunoPET with full-length antibodies to long-lived isotopes such as ^89^Zr (t_1/2_ = 78.4 h) or ^64^Cu (t_1/2_ = 12.7 h), which possess suboptimal physical properties (e.g., beta-branching ratio, R_mean_) compared to more commonly used short-lived isotopes like ^68^Ga (t_1/2_ = 68 min) or ^18^F (t_1/2_ = 109.7 min).^14^ The use of long-lived positron emitters also imposes dose-limiting radiation to healthy organs.^11,15–16^ It has been estimated that the effective dose of [^89^Zr]Zr-labelled mAbs for imaging falls within the range of 0.38 to 0.61 mSv/MBq, and the red marrow dose could potentially reach 0.69 mSv/MBq.^17–23^ This accentuates the need for approaches that reduce radiation exposure. Addressing the limitations associated with the long serum half-life and suboptimal radiation dosimetry of antibody tracers is crucial for maximizing the clinical utility of antibody-based PET imaging. The net results of these physical and pharmacological limitations are poor quality images that take a long time to acquire placing significant burdens on patients and healthcare resources alike. Despite decades of efforts, radiolabeled antibodies have not shown widespread utility nor clinical utilization. Efforts to generate antibody derivatives like single chain Fv, minibodies, diabodies for more favorable kinetics have struggled to achieve the optimal balance between loss of target affinity/specificity and imaging characteristics.^24^ Maintaining the exquisite properties of intact antibodies in a system that permits early, high quality, and high contrast imaging would be a major advance.

Rituximab is a genetically engineered chimeric monoclonal antibody designed to target the CD20 antigen.^25–26^ Initially developed for the treatment of non-Hodgkin’s lymphoma, rituximab has found expanded applications in managing various autoimmune and immune-mediated conditions including rheumatoid arthritis, pemphigus diseases, systemic lupus erythematosus, dermatomyositis, and idiopathic thrombocytopenic purpura.^27^ Given its broad clinical applications, we focused on the development of companion diagnostic PET radioconjugates using rituximab. The serum half-life of rituximab typically ranges from 18 to 22 days.^28^ A recent study assessed [^89^Zr]Zr-labeled rituximab PET as an imaging biomarker to evaluate CD20 targeting in patients with relapsed/refractory diffuse large B-cell lymphoma. The study showed a positive correlation between tumor uptake of [^89^Zr]Zr-rituximab and CD20 expression in tumor biopsies and found the best imaging at approximately 6 days after administration.^29^ Improved techniques for CD20 imaging would be beneficial for new CD20 targeted therapies such as bi-specific antibodies, especially in 3^rd^ and 4^th^ line therapy, where heterogeneity of receptor expression and/or CD20-loss related to prior CD20-directed therapies can impact the likihood of success of CD20-targeted therapy.^30–32^

In this study, we explored the application of biorthogonal chemistry to tackle the challenge associated with the extended blood half-life of rituximab.^33–34^ Our objective is to introduce flexibility into antibody-based imaging, allowing high-contrast imaging at any chosen time following antibody administration. This method, which we call Tetrazine-Knock-Out (TKO), is rooted in transcyclooctene-tetrazine (TCO-Tz) iEDDA chemistry, entails the straightforward modification of antibodies with a TCO linker. This linker was conjugated to different radioisotopes including ^68^Ga, ^125^I (t_1/2_ = 60 days) or ^89^Zr, bound by typical small molecule groups. We tested the ability of antibody tracer to accumulate in tumors from a few hours to a day, and subsequently tested whether an intravenously delivered Tz could efficiently eliminate radioactivity from the circulation and non-target organs (**Fig. 1**). This *in vivo* custom chemical approach enhanced the target-to-background ratio and underscored its potential relevance in clinical settings with specific applications in hematologic malignancies where companion diagnostic imaging may guide therapeutic selection.^35^

**Fig 1.**
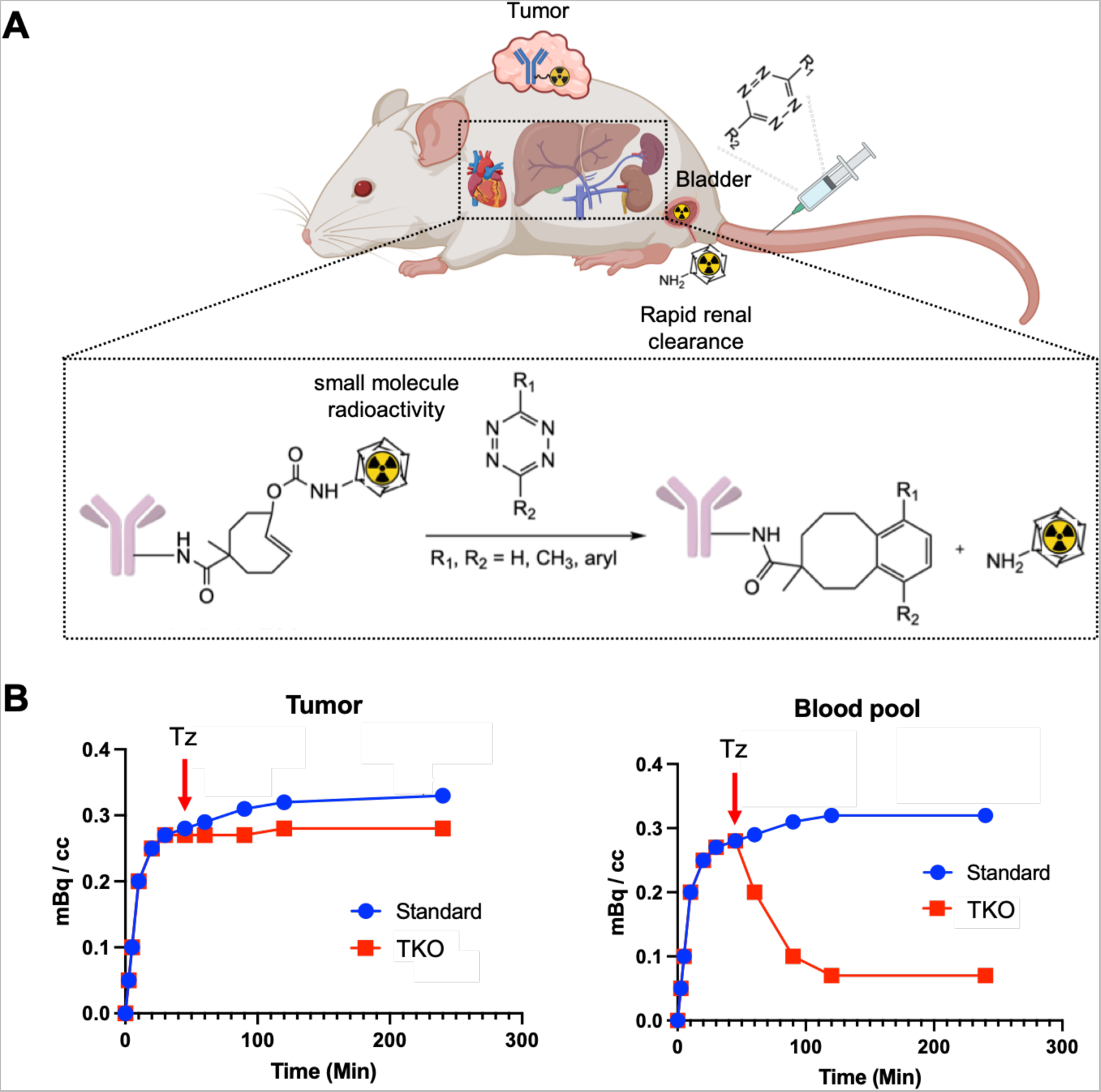
Schematic representations of the Tetrazine-Knock-Out (TKO) approach. **A**) A radioimmunoconjugate contains a trans-cyclooctene linker. The radioactive counterpart of the conjugate can be released by tetrazine (Tz) derivatives from antibodies in circulation and extracellular compartments, followed by renal excretion. Thus, circulating radioactivity outside of the tumor can be effectively eliminated, providing control over the dose in non-target, healthy organs and tissues. **B**) Representative, theoretical curves of expected changes in radioactivity over time in tumor and blood pool.

## Results

### Synthesis of “Tetrazine-Knock Out” (TKO) immunoPET tracers

We synthesized three distinct radio-immunoconjugates, [^125^I]I-AG(Aromatic Group)-TCO-rituximab, [^68^Ga]Ga-NOTA-TCO-rituximab and [^89^Zr]Zr-DFO-TCO-rituximab **(Fig. 2A, Fig. S1A-S3A)**. We used a published method to synthesize the TCO-based small molecule 2 (approximately 100% *trans* based on HPLC/ELSD and ^1^H NMR), beginning with cyclooctadiene 1.^33^ Subsequently, this molecule was coupled with various chelators or small molecules and then combined with rituximab in a sodium bicarbonate buffer at pH 8.5. To ensure the purity of the bioconjugates, we employed centrifugal filtration for purification. The resulting bioconjugates were then stored at 4°C to maintain their stability. Using spectroscopic techniques,^36^ we determined that rituximab is conjugated with approximately 4.5, 2.6, and 3.2 equivalents of AG, NOTA, or DFO, respectively. Subsequently, these bioconjugates were radiolabeled with ^125^I, ^68^Ga and ^89^Zr resulting in corresponding radioimmunoconjugates with a radiochemical purity exceeding 95% as determined by radio-TLC analysis (**Fig. S1-S3**). In the case of AG-TCO-rituximab, we anticipated that radioiodination would preferentially occur with the AG-TCO moiety rather than tyrosine residues on the antibody due to the greater spatial separation between AG-TCO and antibody resulting in an increased exposure of AG-TCO to [^125^I]NaI. A control experiment with unmodified rituximab conducted under the same conditions demonstrated 12% radioiodination of the parent antibody, compared to a radiolabeling efficiency of 86% with AG-TCO-rituximab (**Fig. S1**). To establish the generality of our approach, we applied a similar synthesis strategy to the anti-mouse CD69 antibody H1.2F3,^37^ obtaining [^89^Zr]Zr-DFO-TCO-H1.2F3. We also evaluated the stability of the radiotracers in PBS and FBS, monitoring stability through radio-TLC. All radiotracers exhibited high stability, with no significant degradation observed **(Fig. S1B-S3B**).

**Fig 2.**
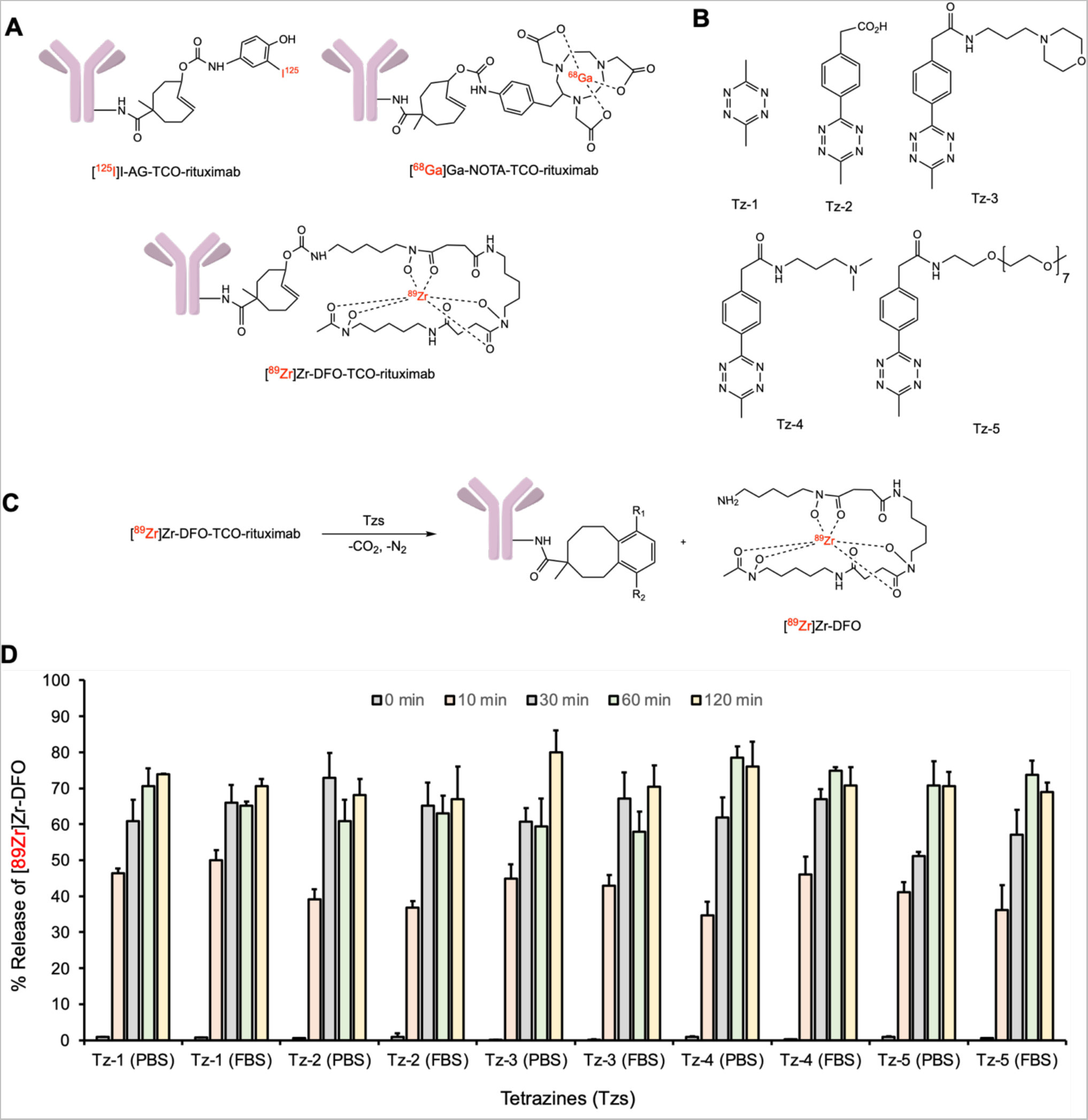
Antibody conjugate structures, Tz derivatives, and in vitro cleavage studies. **A**) TCO-radioimmunoconjugates. Three different radioisotopes with distinct decay properties were used to establish the TKO approach. **B**) Different Tzs used in the study. **C**) Schematic representation of the click reaction to liberate small molecule bearing radioisotope, [^89^Zr]Zr-DFO, from [^89^Zr]Zr-DFO-TCO-rituximab; **D**) In vitro knock out studies. [^89^Zr]Zr-DFO was released from [^89^Zr]Zr-DFO-TCO-rituximab upon reaction with tetrazines (Tz 1-5) in PBS and FBS at 37°C up to 2 h. The progress of the reaction was monitored by radio-TLC (iTLC/citrate buffer (0.1 M, pH 5.0) as the mobile phase). Under these conditions radioimmunoconjugates remained at the origin whereas [^89^Zr]Zr-DFO exhibited an R_f_ of 0.2-0.3. The data represent the mean ± SD (n = 3).

### Tz synthesis

The *in vivo* efficacy of the TKO system is theoretically contingent on factors including 1) the kinetics of the TCO-Tz reaction and 2) the availability of Tz in circulation to facilitate the TKO reaction. Several Tz derivatives were developed to potentially augment the plasma half-life of the clearing agent while maintaining reactivity toward TCO (**Fig. 2B**). The synthesis of Tzs entailed an amine-NHS ester coupling reaction (**Fig. S4**) in DMSO/aqueous Na_2_CO_3_ (pH 8.5) at 37°C, followed by HPLC purification (see supplementary for spectroscopic data). Next, we assessed the *in vitro* toxicity of the Tzs in two different cell types, HEK293T and primary hepatocytes (**Fig. S5**), and the blood retention of all Tzs in a healthy mouse model. No detectable cellular toxicity was observed with any Tz. To check the blood half-lives, equal quantities (0.003 mmol) of Tzs-1-5 were intravenously injected, and blood samples were drawn at 10 minutes post-injection. Serum samples were then analyzed through LCMS and no appreciable amount of any Tz was detected in the blood, suggesting that minor structural differences may not substantially alter the rapid excretion of the Tzs (data not shown).

### Reactivity of Tzs towards cleavable radioimmunoconjugates

We then assessed the reactivity of our five disubstituted tetrazines, Tz-1 to Tz-5, towards [^125^I]I-AG-TCO-rituximab (**Fig. S6**) and [^89^Zr]Zr-DFO-TCO-rituximab (**Fig. 2C-D**). Two distinct leaving groups were used to discern the impact of leaving group characteristics on cleavage efficiency. Incubation of the Tzs with the radioimmunoconjugates was performed in both PBS and FBS (**Fig. 2D, Fig S6B**). The extent of cleavage was monitored qualitatively through radio-TLC. For [^89^Zr]Zr-DFO-TCO-rituximab, cleavage (34-46%) occurred within the initial 10 minutes of the reaction, regardless of the reaction environment (PBS or FBS) or the specific tetrazine used. Cleavage gradually increased to 67-74% over a 2 h duration. In this initial exploration, we used a batch of TCO consisting of approximately 80% *trans* and 20% *cis* isomers. As a result, the reaction proceeded as expected given the purity of the linker. In addition, the Tzs used in our approach exhibited optimal cleavage within the initial 10 minutes of the reaction, after which the reaction rate declined. Commercially available Tz-1, the Tz with lowest molecular weight, exhibited cleavage similar to the other derivatized tetrazines. Despite the anticipated faster release of [^125^I]I-AG from [^125^I]I-AG-TCO-rituximab (**Fig. S6A**) due to the better stability of the released aromatic amines, the cleavage proceeded more slowly compared to [^89^Zr]Zr-DFO-TCO-rituximab (**Fig. 2C**). 3-17% cleavage was observed at 10 minutes, increasing to 46-82% within 2 h. Given the rapid excretion of all Tz derivatives and similar reactivity to TCO-conjugated antibodies, we used commercially available Tz-1 for all of the subsequent experiments.

### Demonstration of TKO strategy in cellular models

Following radiolabeling, we conducted cellular uptake investigations in both CD20^+^ (Raji) and CD20^-^ (K562) cells to confirm the specificity of our radioimmunoconjugates (**Fig 3A**). The observed uptake in Raji cells was 19.93 ± 2.34 %ID/million cells, whereas the uptake in K562 cells was markedly lower at 3.92 ± 3.90 %ID/million cells (n = 3) when exposed to 0.037 MBq of the [^89^Zr]Zr-radioimmunoconjugate. Blocking experiments with 100 μg of parent rituximab resulted in decreased uptake of 2.83 ± 1.90 %ID/million cells. A reduced dose of 0.019 MBq decreased uptake as well 7.67 ± 0.89 %ID/million cells. Comparable trends were observed in cells incubated with [^125^I]I-AG-TCO-rituximab (**Fig. S7A**). To evaluate immunospecificity, we directly radio-iodinated rituximab with ^125^I, using Chloramine T as the oxidizing agent (**Fig. 1C**). The directly radio-iodinated rituximab, [^125^I]I-rituximab, exhibited similar uptake (14.98 ± 0.92 %ID/million cells) to that of [^125^I]I-AG-TCO-rituximab (16.14 ± 0.88 %ID/million cells) (**Fig. S7A**), indicating that the immunoreactivity of the antibody remained unaltered following modification with TCO-AG.

**Fig 3.**
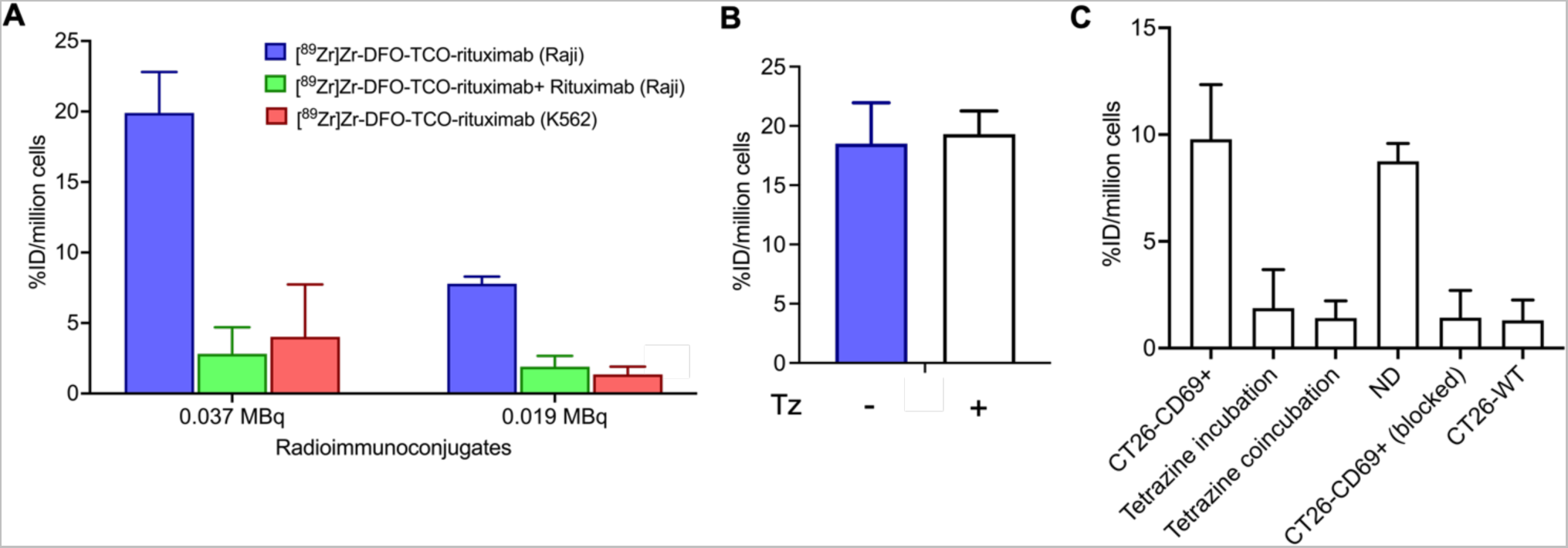
Demonstration of the TKO strategy in cells with rapidly internalizing (rituximab) and slowly internalizing (H1.2F3) antibodies. **A)** Cell uptake study of [^89^Zr]Zr-DFO-TCO-rituximab in Raji (CD20^+^) and K562 cells (CD20^-^). Cells were incubated with the radioimmunoconjugates at 37°C for 2 h in Opti-MEM supplemented with 2% BSA in presence or absence of parent rituximab (100 μg). Uptake was counted in a γ-counter (n = 3). **B)** TKO strategy with an internalizing antibody. Raji cells were first incubated with [^89^Zr]Zr-DFO-TCO-rituximab at 37°C for 2 h in Opti-MEM supplemented with 2% BSA and then incubated at 37°C in presence and absence of Tz-1 for 1 h for selective release of [^89^Zr]Zr-DFO. Cells were then thoroughly washed with 1% DMSO/PBS, centrifuged and counted in a γ-counter (n = 3). No noticeable difference was observed in antibody retention in both Tz-treated and untreated cells. **C)** Demonstration of TKO with a slowly internalizing antibody. CT26-CD69^+^ and CT26-WT cells were incubated with [^89^Zr]Zr-DFO-TCO-H1.2F3 at 25°C for 1 h in Opti-MEM supplemented with 2% BSA in presence and absence of parent H1.2F3 antibody (100 μg). CT26-CD69^+^ cells were then incubated with Tz-1 or with 1% DMSO/Opti-MEM (No-drug, ND) for 1 h for selective release of [^89^Zr]Zr-DFO followed by thorough washing with 1% DMSO/PBS, centrifugation to collect the cell pellets, and counting in a γ-counter (n = 3). Tz coincubation was done with [^89^Zr]Zr-DFO-TCO-H1.2F3 in CT26-CD69^+^ cells under the same condition. Cells were thoroughly washed with 1% DMSO/PBS before counting in a γ-counter (n = 3).

To confirm that Tz treatment would not decrease radioimmunoconjugate uptake in antigen-positive target tissues, Raji cells were additionally incubated with 3,6-dimethyl-1,2,4,5-tetrazine (Tz-1) for 1 h. No significant drop in uptake was observed (18.46 ± 3.89 %ID/million cells vs. 19.45 ± 1.90%ID/million cells) in tetrazine-treated cells (**Fig. 3B, Fig. S7B**). This provides crucial evidence that after antigen binding and internalization, radioimmunoconjugates are not susceptible to substantial reaction with Tzs, allowing for the maintenance of accumulated signal in target cells.

While rituximab is a rapidly internalizing antibody^38–39^, we also tested a slowly internalizing antibody, H1.2F3, targeting the canonical marker for immune activation CD69^37,40–42^. These studies used CT26 mouse colon carcinoma cells transduced to express CD69 (they do not endogenously express CD69, **Fig. S8**). High radiotracer uptake was observed within 1h in CT26-CD69^+^ cells (9.83 ± 2.33% ID/million cells) when incubated with 0.037 MBq of [^89^Zr]Zr-DFO-TCO-H1.2F3 compared to WT cells (1.29 ± 1.09% ID/million cells) (**Fig. 3C**). A substantial drop in tracer uptake in CD69^+^ cells was noted when blocking with 100 μM non-radiolabeled H1.2F3 (1.44 ± 1.22% ID/million cells). This data suggests retained specificity and binding affinity of the TCO-linked antibody tracer in cell culture. Sequential incubation of CT26-CD69^+^ cells initially for 1 h, followed by treating with Tz-1 for another 1 h, resulted in a substantial drop in cellular uptake (1.63 ± 1.94% ID/million cells), which suggests extracellular availability of the TCO to the Tz molecule for cleavage. Co-incubation of CT26-CD69^+^ cells with a mixture of [^89^Zr]Zr-DFO-TCO-H1.2F3 and Tz-1 resulted in no considerable uptake (1.34 ± 0.78% ID/million cells). Thus, the similarly low tracer retention in the sequential Tz incubation and Tz co-incubation groups is likely due to the slow internalization of the antibody into the cells. Similar observations were also noted when the same study was conducted with [^125^I]I-AG-TCO-H1.2F3 (**Fig. S9**). These studies concluded that the radioactivity remaining outside the cells can be eliminated by the introduction of Tzs, emphasizing that the antibody internalization is a factor influencing the timing of Tz administration, at least in *in vitro* models.

### In vivo demonstration of TKO in a healthy mouse model

We intravenously administered [^125^I]I-AG-TCO-rituximab to healthy female mice and waited for 3 h, followed by intravenous administration of Tz or vehicle control (**Fig. 4A**). An ex-vivo biodistribution study was performed after 1 h of Tz/vehicle injection. Upon interaction with Tz, the TCO conjugate released the radioactive leaving group ([^125^I]I-AG, **Fig. S6A**), resulting in a reduction in blood radioactivity from 25.16 ± 1.23 %ID/g to 8.28 ± 0.33%ID/g within 1 h of Tz administration. Mice treated with the vehicle did not show [^125^I]I-AG in urine, while the liberated [^125^I]I-AG was found in the urine of Tz-treated mice by radio-HPLC (**Fig. 4B**). This suggests that the released [^125^I]I-AG rapidly passes through the kidneys and accumulates in bladder for elimination. *In vivo* deiodination resulted in the presence of free ^125^I in the radiochromatogram of both Tz and vehicle treated groups, which is a common occurance with iodine-based radiotracers.

**Fig. 4.**
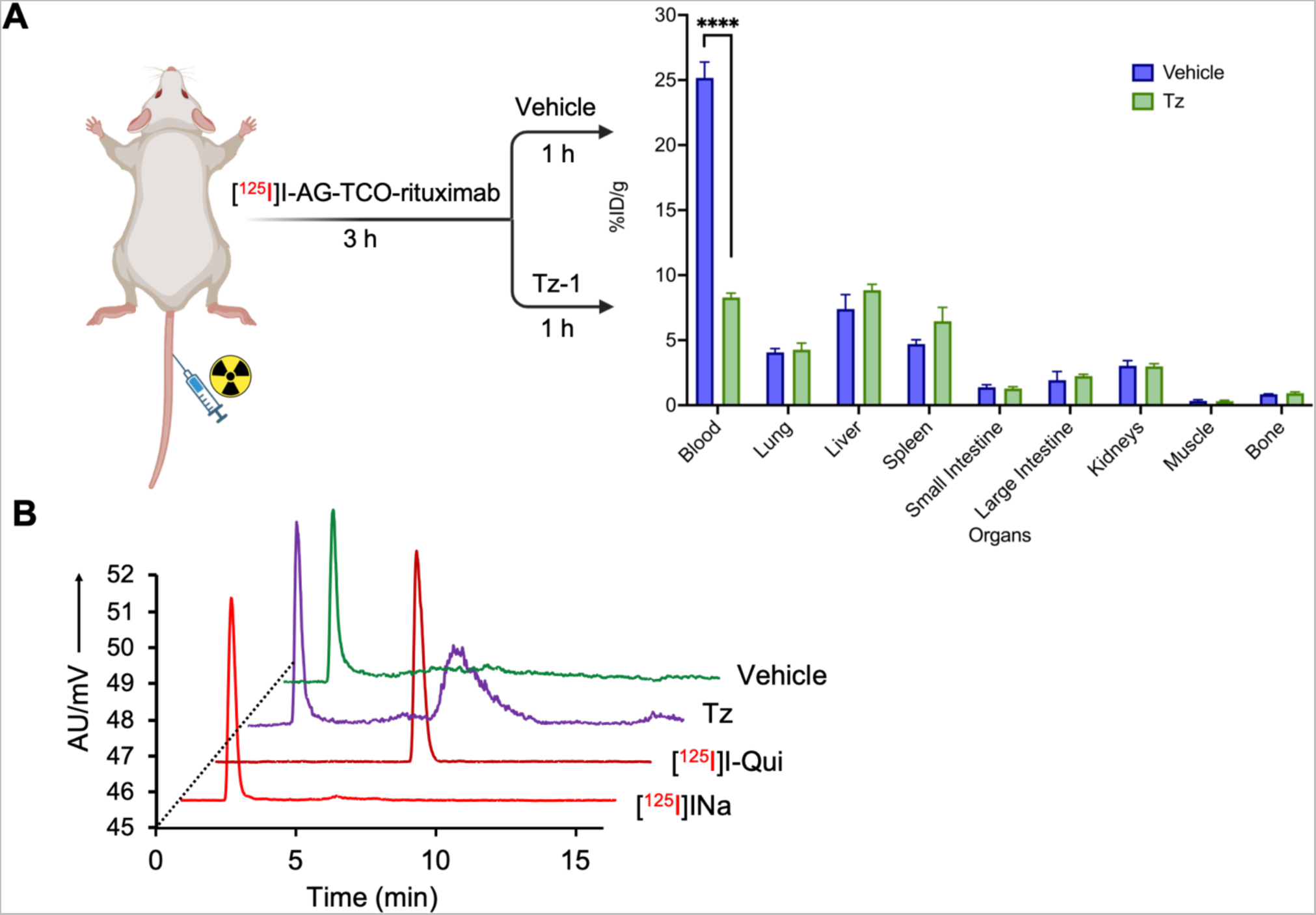
Healthy mouse biodistribution and demonstration of the TKO strategy in Balb/c mice using [^125^I]I-AG-rituximab and Tz-1. **A**) Schematic representation illustrating the injection of [^125^I]I-AG- rituximab, followed by vehicle (5% EtOH/PBS) or Tz administration and subsequent biodistribution (n = 3); **B**) Urine analysis. Urine was collected from both vehicle and tetrazine-treated groups, and the presence or absence of [^125^I]I-AG was confirmed by radio-HPLC.

### In vivo demonstration in tumor xenografts

Next, we tested the TKO strategy in xenograft models with a relatively short-lived isotope, ^68^Ga, to test whether Tz clearance of the radioactive leaving group would result in improved imaging within a few hours of antibody administration. For this study, we expressed CD20 on I45 mesothelioma cells (**Fig. S10**). I45-CD20^+^ tumors were then grown for approximately 2 weeks prior to administration of 18.5 MBq of [^68^Ga]Ga-NOTA-TCO-rituximab. Radioactivity was then allowed to accumulate in tumors for 2 h. At this point, Tz-1 (50 mg/kg) or vehicle (5% EtOH/PBS) was injected i.v. (n = 3), followed by a 1 h wait before doing PET/BioD, following a procedure described in **Fig. 5A**. *Ex vivo* biodistribution (**Fig. 5B**) demonstrated a significant decrease in signal in the blood after the introduction of Tz in animals (48.75 ± 10.47 %ID/g vs. 20.40 ± 4.62 %ID/g for vehicle and Tz groups respectively), as well as in liver (36.71 ± 6.42 %ID/g vs. 23.08 ± 7.21 %ID/g for vehicle and Tz groups respectively) and other background organs like heart, spleen, bone, and lymph node. A representative image of Tz and vehicle-treated mice is shown in **Fig. S11A** and **Fig. S11B** with corresponding SUV_max_ ratios in **Fig. 5C**. Once Tz reacted with the radioimmunoconjugate, the liberated small molecule, [^68^Ga]Ga-NOTA, accumulated in the bladder, which was observed using dynamic PET imaging (**Fig. 5D**). Bladder uptake began increasing immediately after Tz administration and reached saturation within 10 min. In this particular tumor model, a decrease in tumor uptake was observed in Tz treated groups (8.51 ± 2.83 %ID/g vs. 4.21 ± 0.67 %ID/g for vehicle and Tz groups respectively). To explain this decrease in tumor uptake, an *in vitro* experiment was conducted, with I45- CD20^+^ cells incubated with [^68^Ga]Ga-NOTA-TCO-rituximab, followed by Tz-1. A decrease in cellular uptake was seen (11.85 ± 0.09 %ID/million cells and 6.17 ± 0.15 %ID/million cells before and after incubation with Tz, **Fig. S12**), mirroring the *in vivo* findings. This is presumed to be related to the slower rate of internalization of the radioimmunoconjugates into the engineered mesothelioma cells, which may not have the same cellular signaling and protein processing machinery for CD20 internalization compared to B lymphocytes. This suggests that, in general, internalization kinetics must be balanced with incubation time. Notably, a complete clearance of radioactivity from the tumor was observed when a canonical non-internalizing CD20 antibody like obinutuzumab was used (**Fig. S13**).

**Fig 5.**
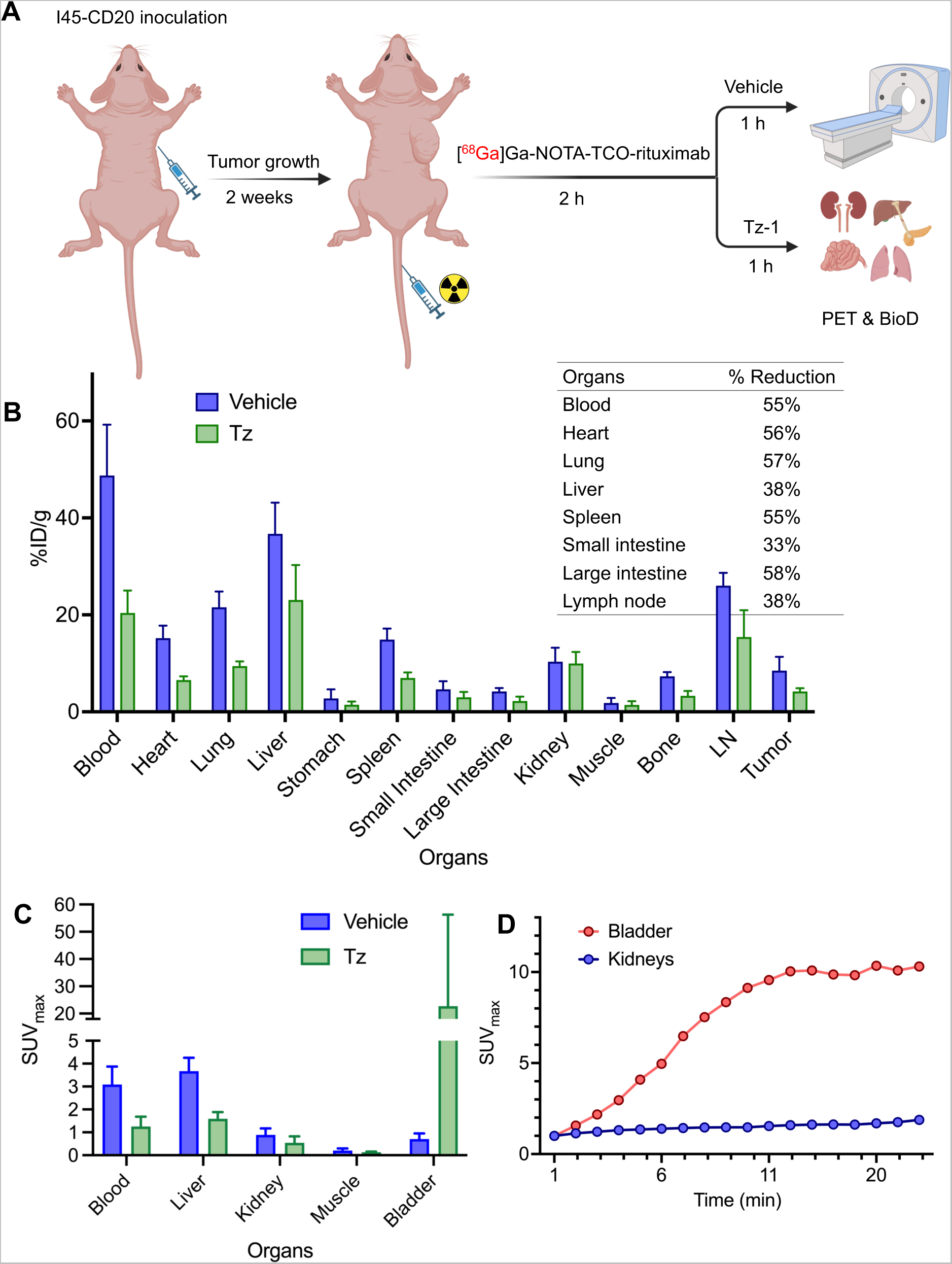
TKO studies with I45-CD20 xenografts at early time points. Female nu/nu mice bearing I45- CD20^+^ xenografts were administered with [^68^Ga]Ga-NOTA-TCO-rituximab and imaging and biodistribution studies were conducted. **A**) Schematic representation illustrating injection of [^68^Ga]Ga-NOTA-TCO-rituximab, followed by Tz/vehicle (5% EtOH/PBS) administration. Animals were scanned in a microPET scanner and euthanized for subsequent biodistribution studies. **B**) Biodistribution of Tz and vehicle treated groups (n = 3). **C**) SUV_max_ for blood, liver, kidney, muscle and bladder for vehicle and Tz treated groups. **D**) SUV_max_ for bladder and kidneys after Tz administration recorded in a dynamic PET scan. The radioactive cleaved molecule quickly passed through the kidneys and rapidly accumulated in the bladder.

Next, we used a clinically-relevant model of B cell lymphoma to investigate the TKO approach using ^89^Zr.^29^ Nude mice bearing Raji tumors, grown for approximately 3 weeks, were administered 7.4 MBq of [^89^Zr]Zr-DFO-TCO-rituximab. After 24 h, intravenous injection of Tz-1 (50 mg/kg) or a vehicle (5% EtOH/PBS) was performed, followed by imaging and biodistribution (**Fig. 6A**). Representative PET images (**Fig. 6B** and **C**) showed a significant reduction in background signal across critical organs within 1 h after Tz administration. PET imaging at 8 and 24 h using an IgG isotype control antibody conjugated with [^89^Zr]Zr-DFO-TCO (**Fig. 6D**) showed no tumor visualization, confirming the tumor specificity of the CD20-targeted radioimmunoconjugate *in vivo*. **Fig. 6E** describes the corresponding SUV_max_ ratios of tumor-to-blood and tumor-to-liver in Tz and vehicle treated groups. Biodistribution studies (**Fig. 6F**) corroborate these findings, revealing a marked decrease in activity across various organs with retention of radioactivity in tumor tissues. This resulted in maximum retention of radioactivity in tumor tissues. There was a substantial reduction (56%) in blood radioactivity (17.66 ± 1.20 %ID/g vs. 7.78 ± 0.59 %ID/g for vehicle and Tz groups respectively), and a decline (27%) in liver uptake upon Tz treatment (8.61 ± 0.82 %ID/g vs. 6.35 ± 1.58 %ID/g for vehicle and Tz groups respectively). A decrease in activity was also seen in lymph node (62%), small and large intestine (45% and 23%). Tumor-associated activity remained unaffected (9.93 ± 1.28 %ID/g vs. 11.81 ± 2.29 %ID/g for vehicle and Tz groups respectively), yielding high-contrast visualization of the tumor in the Tz treated group. Greater than two-fold increases in tumor-to-blood (2.2 times) and tumor-to-liver (2.3 times) ratios were observed (**Fig. 6E**) after 1 h of Tz introduction. Of note, in the vehicle group, tumor to liver ratio was 1.15, making a hepatic tumor difficult or impossible to discern on imaging whereas in the treated group the tumor to liver ratio was 1.86, a ratio easily detectable with PET imaging. Notably, typical [^89^Zr]Zr-DFO-rituximab imaging occurs 3-6 days post-administration in patients, and significant background activity persists in blood pool and liver regions.^30^ To further confirm that the liberated small molecule, [^89^Zr]Zr-DFO, was cleared quickly and did not contribute significantly to imaging signal, we injected [^89^Zr]Zr-DFO in healthy Balb/c mice and performed biodistribution analyses at 1, 3 and 24 h post-injection (**Fig. S14**). At 1 h post-injection, minimal tracer retention was detected in critical organs including blood, liver, spleen, intestine, and kidneys. Kidneys showed the highest radioactivity accumulation (2.13 ± 1.08 %ID/g) followed by liver (2.06 ± 0.01 %ID/g). This low uptake confirmed that [^89^Zr]Zr-DFO is quickly excreted after being knocked out from [^89^Zr]Zr-DFO-TCO-rituximab (**Fig. 2C**), leaving little background signal.

**Fig 6.**
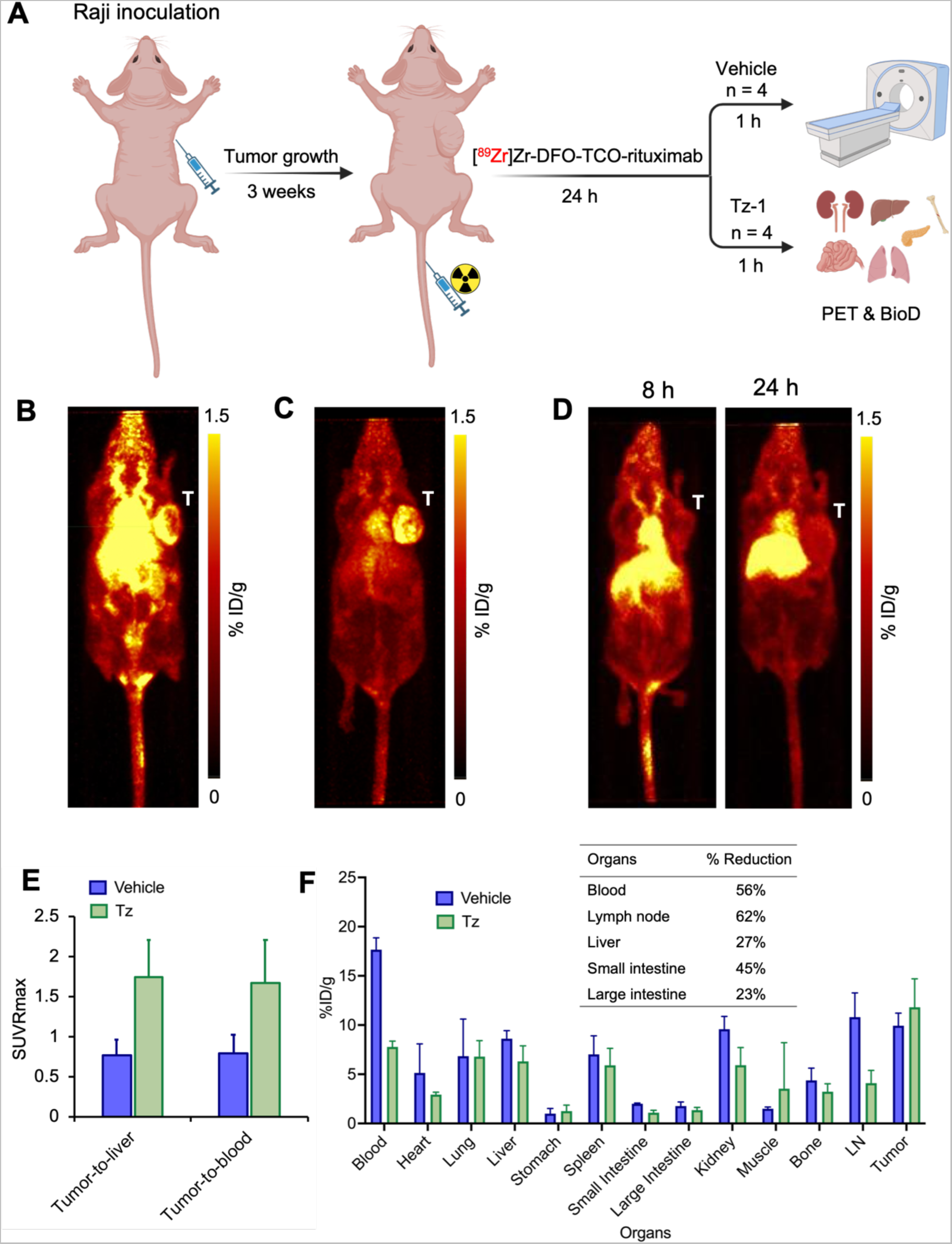
TKO studies with Raji xenografts. *In vivo* demonstration of the TKO strategy in female nu/nu mice bearing Raji xenografts using [^89^Zr]Zr-DFO-TCO-rituximab. **A**) Schematic representation illustrating injection of [^89^Zr]Zr-DFO-TCO-rituximab, followed by Tz/vehicle (5% EtOH/PBS) administration. Animals were scanned in a microPET scanner and euthanized for subsequent biodistribution studies. **B**) Representative PET image for the vehicle-treated group. **C**) Representative PET image for the Tz-treated group. **D**) Representative PET image for isotype control antibody [^89^Zr]Zr- DFO-TCO-IgG1 at 8 and 24 h post-injection. T indicates tumor. **E**) SUVR_max_ for tumor-to-liver and tumor- to-blood for Tz and vehicle treated groups (n = 4). **F**) Biodistribution of Tz and vehicle treated groups (n = 4).

In both *in vivo* studies, the TKO strategy showed a similar 50-60% decrease in off-target radioactivity following a single dose of Tz injection. Our Tz, with a small molecular weight (∼110 g/mol) and very short blood half-life, retained its reactivity when modified with various short linkers (**Fig. 2D**), though the potential effects of multiple or high molecular weight Tzs on further reducing background radiation remain unexplored.

## Discussion

Conjugating radioactivity to biologic molecules such as antibodies has been an important area of exploration for molecular imaging and radioimmunotherapy (RIT). Typically, the long circulating blood half-lives of radioimmunoconjugates necessitates the injection of activities 10-fold lower than those used for small molecule radiotracers like [^18^F]FDG with the optimal imaging window several days after injection. Given these limitations and despite 30 years of research, antibody-based PET imaging agents are not used widely. It is notable that the ZIRCON trial, using [^89^Zr]-DFO-girentuximab to image the renal cell carcinoma antigen carbonic anhydrase IX, has completed late phase clinical trials and has regulatory approval,^43^ suggesting continued efforts to use radioimmunoconjugates in the clinic. Moreover, dose limitations make RIT complicated and sometimes untenable when using therapeutic isotopes, with a narrow therapeutic window and potential damage to healthy organs, including bone marrow. Zevalin and Bexxar, despite FDA approval, are rarely used or marketed and solid tumor RIT has failed to show sufficient activity in clinical trials (NCT01956812). Hence, there exists a clear need to improve upon standard approaches for full length antibody nuclear imaging and therapy. Our goal was to explore custom chemical approaches that balance the duration of antibody-isotope conjugate in the body to ensure sufficient accumulation in tumors, while increasing the clearance of radioactivity from non-target tissues.

The reaction between TCO and Tz is the fastest *in vivo* biorthogonal reaction,^44^ and iEDDA chemistry is being used for pre-targeted nuclear imaging^42^ and *in vivo* drug delivery.^33–34^ Pre-targeted antibody tumor imaging is an emerging technique in cancer diagnostics and therapeutics, and can enhance the tumor-to-background ratio. This method also uses a two-step procedure: initially, a non-radioactive antibody, often modified with TCO, is administered, selectively binding to cancer-associated markers or antigens. Following this step, there is a waiting period of 1-3 days (or longer) to eliminate sufficient unbound antibody from circulation; or sometimes an antibody clearing agent is also used. Subsequently, a radiolabeled Tz is introduced to react with the TCOs on the previously administered antibody in the tumor microenvironment. This sequential process facilitates the accumulation of the radiolabeled probe selectively at the tumor site and allows the radioactivity to be conjugated to a small molecule.^45^

The TKO approach diverges from these pre-targeting antibody imaging methods in that the goal is to use the click reaction not to attach the radioactivity at a later time in tumor tissues, but to knock out the radioactivity from normal tissues, especially the blood, at an opportune time for either diagnostic or therapeutic efficacy. This borrows concepts from click-to-release chemotherapies and an early attempt has been made to try release chemistry using Staudinger ligation for radioactivity, but relatively small improvements were observed.^46^ In contrast to alternatives, the TKO strategy offers several advantages. The TKO reaction occurs primarily in the bloodstream, where Tz concentration is highest after administration, rather than at the tumor site, at a favorable stoichiometry for efficient *in vivo* reaction. The Tz-TCO reaction rate is incredibly fast, as evidenced by our dynamic scan showing rapid bladder accumulation of radioactivity (**Fig. 5B**). Finally, there is no rigid need for a prolonged waiting period to clear circulating antibodies as is the case with pre-targeting approaches. A recent report by Vlastara et al. showed an increase in tumor-to-blood ratio with a [^89^Zr]Zr-trastuzumab radioimmunoconjugate using [^89^Zr]Zr-DFO as the leaving group.^47^ Our study more broadly evaluates different radioisotopes, leaving groups, cell lines, antibodies, and *in vivo* characteristics of the system.

The rate of antibody internalization into target cells has an important role in the TKO approach in this first, most checmically straightforward instantiation. While slowly internalizing or non-internalizing antibodies may be more difficult to employ with the TKO approach, there are a substantial number of FDA-approved robustly internalizing antibodies like trastuzumab (HER2), cetuximab (EGFR), bevacizumab (VEGF), pertuzumab (HER2), daratumumab (CD38), brentuximab (CD30), ipilimumab (CTLA-4), atezolizumab (PD-L1), and several antibody drug conjugates (ADCs) or antibodies in clinical trials.^48–52^ These antibodies underscore the potential generality of our approach. Furthermore, the modular nature of the TKO strategy allows its application to virtually any antibody, antibody fragment, or macromolecule (including nanoparticles) conjugated to any radioisotope, including isotopes like ^18^F, which have a favorable half-life for single day injection and imaging as well as ideal physical properties. Future studies will explore the potential of the TKO platform for theranostic pairs with ^18^F and ^131^I or ^211^At.

To demonstrate the TKO approach, we selected rituximab as a prototypical internalizing antibody with significant clinical utility and a long serum half-life.^28^ Once rituximab binds to CD20 on target cells, it is internalized and shielded from the TCO-Tz reaction in B cells. Cells lacking CD20 (i.e., most cells in the body), do not internalize the antibody. Any antibodies remaining in circulation or on the cell surface could react with Tzs and release the radioactivity for rapid clearance (**Fig. 3B and S7**). This difference in susceptibility to the TCO-Tz reaction was confirmed *in vivo* in Raji xenograft models (**Fig. 6B-C**). By transitioning the radionuclide from a slowly clearing radioimmunoconjugate into a rapidly clearing small molecule, the TKO approach effectively reduces radiation exposure to these non-target organs while achieving sufficient target to background for imaging days earlier than the conventional approach. Similar observations were also noticed when TKO was conducted with [^68^Ga]Ga-NOTA-TCO-rituximab in I45-CD20^+^ xenografts. However, in this case, a decrease in tumor radioactivity (**Fig. 5B**) was also observed, likely because the engineered cells require more time for internalization. Extending the circulating time of the antibody could be a potential solution to address this issue.

The success of our approach also relies on the serum half-life of Tzs and their rapid reaction with TCOs. All the Tzs (**Fig. 2B**) we used in our study showed equal reactivity in *in vitro* studies in FBS (**Fig. 2D and S6B**) and were not detectable in the blood within 10 minutes of introduction using standard techniques. Tz-1 was used for *in vivo* studies (**Fig. 4-6**) and is effectively the most minimal Tz derivative; nonetheless, we observed 50-60% reduction in background signal with a single dose. Administering Tzs with higher molecular weight, longer chain polyethylene glycol (PEG) groups, or in multiple doses could potentially lead to greater decreases in background radiation.

In summary, we show that the TKO approach can improve the imaging contrast of radioimmunoconjugates at various time points. The TKO approach has significant promise for advancing cancer diagnosis, optimizing antibody imaging agents in predictive biomarker imaging, and improving the safety and dosimetry of radiotherapeutic and theranostics.

## Materials and Methods

### General procedures

Chemicals were sourced from commercial suppliers at high purity. Monitoring of reactions involving non-radioactive compounds was performed through thin-layer chromatography (TLC) analysis employing precoated silica gel 60 F254 plates. Column chromatography utilized silica gel (100–200 mesh), while the final purification of compounds was accomplished using an Agilent 1260 Infinity II High-Performance Liquid Chromatography (HPLC) system. Nuclear magnetic resonance (NMR) analyses were performed on either a Bruker NEO400 spectrometer equipped with a 5mm BBFO IProbe (^1^H = 400 MHz and ^13^C = 100 MHz) or a Bruker NEO600 spectrometer with a Prodigy BBO cryoprobe (^1^H = 600 MHz and ^13^C = 150 MHz). Chemical shifts are reported in parts per million (ppm), and coupling constants are expressed in Hz, with abbreviations such as s (singlet), d (doublet), dd (doublet of doublets), t (triplet), q (quartet), and m (multiplet) used to denote splitting patterns. For nominal mass accuracy Liquid Chromatography-Mass Spectrometry (LCMS) data, we employed a Waters Acquity UPLC system equipped with a Waters TUV detector (254 nm) and a Waters SQD single quadrupole mass analyzer with electrospray ionization. The LC gradient included a 30-second hold at 95:5 (water:acetonitrile, 0.1% v/v formic acid), followed by a 2-minute gradient to 5:95, and concluding with a 30-second hold. The column utilized was an Acquity UPLC HSS C18, 1.7um, 2.1x 50 mm. [^89^Zr]Zr-oxalate was procured from 3D Imaging, USA. [^125^I]NaI was purchased from Perkin Elmer (0.1 M NaOH (pH 12-14), reductant-free, specific activity ∼17 Ci (629 GBq)/mg). In-house generation of [^68^Ga]GaCl_3_ took place at the University of Pennsylvania Cyclotron & Radiochemistry Facility from ^68^Ge/^68^Ga generators eluted with 0.1 N HCl. TLC plates were either scanned using a radio-TLC scanner or divided into equal pieces and counted in a gamma counter.

### Synthesis of NOTA-TCO-rituximab (4)

Compound 2 was synthesized according to published methods.^30^ To a solution of compound 2 (5 mg, 0.01 mmol) in DMSO (100 μL), NaHCO_3_ was added dropwise until pH of the reaction mixture reached ∼8.5. To this mixture NOTA-Bn-NH_2_ (5 mg, 0.009 mmol, Macrocyclics) was added and the mixture was incubated at 37°C for 4 h. It was then diluted with water (2.5 mL) and passed through a RP-HPLC column. The volatiles were removed using a SpeedVac concentrator (Fisher Scientific), and the light-yellow solid (0.45 mg) was redissolved in dry DMSO (0.45 mL). Rituximab (1.7 mg in 100 μL PBS, pH = 7.4) was then added to the 10 μL of the above mixture and the total volume was adjusted to 0.5 mL with PBS. A few drops of Na_2_CO_3_ (0.1 M in water) were then added in the reaction mixture to adjust the pH to 8-9. The mixture was incubated at 37°C while shaking at 400 rpm for 4 h in complete darkness before concentration using an Amicon YM-10 centrifugal filter (washed twice with 1% DMSO/H_2_O and then with H_2_O).

### Synthesis of [^68^Ga]Ga-NOTA-TCO-rituximab

NOTA-TCO-rituximab (100 μg, 10 μL water) in 2 M sodium acetate buffer (100 μL, pH 5.5) was placed in a screw tube. Carrier-free [^68^Ga]GaCl_3_ (374 MBq) in 0.1 N HCl (1.1 mL) was added to this solution (final pH of the reaction mixture was ∼5.5). The reaction mixture was incubated for 30 min at 37°C shaking at 400 rpm. The progress of reaction was tracked using radio-TLC with Varian ITLC-SA strips and 50 mM EDTA (pH 4.5) as the mobile phase. In this system, unbound ^68^Ga forms a complex with EDTA, migrating to the solvent front (R_f_ = 1), while [^68^Ga]Ga-NOTA-TCO-rituximab remained at the origin (R_f_ = 0). Purification was achieved through a Cytiva PD-10 column, resulting in the isolation of pure [^68^Ga]Ga-NOTA-TCO-rituximab (74 MBq) with a radiochemical purity exceeding 95% (specific activity 0.74 MBq/µg).

### Synthesis of DFO-TCO-rituximab (4)

To the solution of compound 2 (5 mg, 0.01 mmol) in DMSO (500 μL), NaHCO_3_ was added dropwise until pH of the reaction mixture reached ∼8.5. To this mixture DFO-NH_2_ mesylate (7 mg, 0.01 mmol, Cayman) was added and the mixture was incubated at 37 °C for 4 h. It was then diluted with water (2.5 mL) and passed through a RP-HPLC column. The volatiles were removed using a SpeedVac concentrator (Fisher Scientific) and the yellow gel (1.1 mg) obtained was redissolved in dry DMSO (0.11 mL). Rituximab (1.7 mg in 100 μL PBS, pH = 7.4) was then added to the 10 μL of the above mixture and the total volume was adjusted to 0.5 mL with PBS. A few drops of Na_2_CO_3_ (0.1 M in water) were then added in the reaction mixture to adjust the pH to 8-9. The mixture was then shaken at 400 rpm for 4 h at 37 °C in complete darkness and concentrated using a YM-10 Amicon centrifugal filter (washed twice with 1% DMSO/H_2_O and then with H_2_O).

### Synthesis of [^89^Zr]Zr-DFO-TCO-rituximab

Radiolabeling of DFO-TCO-rituximab with ^89^Zr involved the reaction of 100 µg (10 µL in water) of DFO-TCO-rituximab with carrier free [^89^Zr]Zr-oxalate (81 MBq) in 200 µL of 1 M HEPES (pH 7) buffer. The reaction mixture was incubated while shaking at 400 rpm for 30 minutes at 37°C. The progress of [^89^Zr]Zr-DFO-TCO-rituximab formation was tracked using radio-TLC with Varian ITLC-SA strips and 0.1 M EDTA (pH 5) as the mobile phase. In this system, unbound [^89^Zr]Zr forms a complex with EDTA, migrating to the solvent front (R_f_ = 1), while [^89^Zr]Zr-DFO-TCO-rituximab remained at the origin (R_f_ = 0). Purification was achieved through a Cytiva PD-10 column, resulting in the isolation of pure [^89^Zr]Zr-DFO-TCO-rituximab (74 MBq) with a radiochemical purity exceeding 95% (specific activity 0.74 MBq/ µg).

### Synthesis of tetrazine Tz-3

In a DMSO solution (0.5 mL) containing 3-morpholinopropan-1-amine (4, 10 mg, 0.07 mmol), 1 M Na_2_CO_3_ was added dropwise until the pH of the reaction mixture reached approximately 8.5. Subsequently, 2,5-dioxopyrrolidin-1-yl 2-(4-(6-methyl-1,2,4,5-tetrazin-3-yl)phenyl)acetate (5, 23 mg, 0.07 mmol) was added, and the mixture was heated at 37°C for 2 h, during which LCMS analysis confirmed the formation of the desired compound. The solvent was then diluted with water (DMSO:water = 1:4) and subjected to purification through preparative reverse-phase HPLC chromatography (Agilent 1260 Infinity II HPLC). The chromatographic separation was carried out on an Agilent Prep-C18 column (250 mm × 21.2 mm × 10 μm) using a linear gradient (5% to 95% in 20 min, flow-rate of 20 mL/min) of solvent B (0.1% TFA in ACN, v/v) in solvent A (0.1% TFA in H2O, v/v). UV detection was set at 254 and 280 nm. The volatiles were removed using a SpeedVac Concentrator (Fisher Scientific), and the resulting product, Tz-3, was obtained as a bright-pink solid (8 mg, 32%) after lyophilization. ^1^H NMR (400 MHz, DMSO) δ 8.41 (d, *J* = 8.5 Hz, 2H), 8.13 (t, *J* = 5.6 Hz, 1H), 7.54 (d, *J* = 8.5 Hz, 2H), 3.60 – 3.49 (m, 6H), 3.15 – 3.06 (m, 2H), 3.00 (s, 3H), 2.36 – 2.20 (m, 6H), 1.57 (p, *J* = 7.1 Hz, 2H) ppm; ^13^C NMR (100 MHz, DMSO) δ 169.82, 167.52, 163.71, 141.82, 130.52, 130.49, 127.78, 66.67, 56.26, 53.80, 42.84, 40.64, 40.43, 40.22, 40.02, 39.81, 39.60, 39.39, 37.52, 26.48, 21.29 ppm; LCMS: 358.3 [M+2H]^+^; HRMS: calcd for C_18_H_24_N_6_O_2_ [M+H]^+^ 357.2039, found 357.2040.

### Synthesis of tetrazine Tz-4

In a DMSO solution (0.5 mL) containing N,N-dimethylpropane-1,3-diamine (6, 12 mg, 0.12 mmol), 1 M Na_2_CO_3_ was added dropwise until the pH of the reaction mixture reached approximately 8.5. Subsequently, 2,5-dioxopyrrolidin-1-yl 2-(4-(6-methyl-1,2,4,5-tetrazin-3-yl)phenyl)acetate (5, 38 mg, 0.12 mmol) was added, and the mixture was heated at 37°C for 2 h, during which LCMS analysis confirmed the formation of the desired compound. The solvent was then diluted with water (DMSO:water = 1:4) and subjected to purification through preparative reverse-phase HPLC chromatography (Agilent 1260 Infinity II HPLC). The chromatographic separation was carried out on an Agilent Prep-C18 column (250 mm × 21.2 mm × 10 μm) using a linear gradient (5% to 95% in 20 min, flow-rate of 20 mL/min) of solvent B (0.1% TFA in ACN, v/v) in solvent A (0.1% TFA in H2O, v/v). UV detection was set at 254 and 280 nm. The volatiles were removed using a SpeedVac concentrator (Fisher Scientific), and the resulting product, Tz-4, was obtained as a bright-pink solid (3.2 mg, 8%) after lyophilization. ^1^H NMR (600 MHz, DMSO-d_6_) δ 8.42 (d, *J* = 8.4 Hz, 2H), 8.33 (t, *J* = 5.8 Hz, 1H), 7.56 (d, *J* = 8.6 Hz, 2H), 3.58 (s, 2H), 3.15 (q, *J* = 6.7 Hz, 2H), 3.06 – 3.01 (m, 2H), 3.00 (s, 3H), 2.76 (d, *J* = 4.9 Hz, 6H), 1.83 – 1.74 (m, 2H) ppm; ^13^C NMR (100 MHz, DMSO-d_6_) δ 170.47, 167.56, 163.70, 141.50, 130.61, 127.84, 55.15, 42.77, 40.63, 40.42, 40.22, 40.01, 39.80, 39.59, 39.38, 36.27, 24.88, 21.29 ppm; LCMS: 316.4 [M+2H]^+^; HRMS: calcd for C_16_H_22_N_6_O_2_ [M+H]^+^ 315.1933, found 315.1988

### Synthesis of tetrazine Tz-5

In a DMSO solution (0.5 mL) containing 2,5,8,11,14,17,20,23-octaoxapentacosan-25-amine (7, 8 mg, 0.023 mmol), 1 M Na_2_CO_3_ was added dropwise until the pH of the reaction mixture reached approximately 8.5. Subsequently, 2,5-dioxopyrrolidin-1-yl 2-(4-(6-methyl-1,2,4,5-tetrazin-3-yl)phenyl)acetate (5, 7.7 mg, 0.023 mmol) was added, and the mixture was heated at 37°C for 2 h, during which LCMS analysis confirmed the formation of the desired compound. The solvent was then diluted with water (DMSO:water = 1:4) and subjected to purification through preparative reverse-phase HPLC chromatography (Agilent 1260 Infinity II HPLC). The chromatographic separation was carried out on an Agilent Prep-C18 column (250 mm × 21.2 mm × 10 μm) using a linear gradient (5% to 95% in 20 min, flow-rate of 20 mL/min) of solvent B (0.1% TFA in ACN, v/v) in solvent A (0.1% TFA in H_2_O, v/v). UV detection was set at 254 and 280 nm. The volatiles were removed using a SpeedVac concentrator (Fisher Scientific), and the resulting product, Tz-5, was obtained as a bright-pink solid (6 mg, 48%) after lyophilization. ^1^H NMR (400 MHz, DMSO-d_6_) δ 8.41 (d, *J* = 8.5 Hz, 2H), 8.23 (t, *J* = 5.7 Hz, 1H), 7.55 (d, *J* = 8.6 Hz, 2H), 3.58 (s, 2H), 3.56 – 3.47 (m, 26H), 3.47 – 3.39 (m, 4H), 3.24 (s, 5H), 3.00 (s, 3H) ppm; ^13^C NMR (100 MHz, DMSO-d_6_) δ 170.07, 167.52, 163.71, 141.76, 130.54, 130.48, 127.76, 71.75, 70.25, 70.21, 70.09, 70.05, 69.54, 58.51, 42.64, 40.52, 40.42, 40.22, 40.01, 39.80, 39.59, 39.38, 21.28 ppm; LCMS: 597.9 [M+2H]^+^; HRMS: calcd for C_26_H_45_N_5_O_9_ [M+H]^+^ 596.3296, found 596.3302.

### In vitro cleavage study using Tz-1-5

The radioimmunoconjugate [^89^Zr]Zr-DFO-TCO-rituximab (1.85 MBq) was incubated with Tz-1-5 (1 mg, 5% EtOH) in either PBS (pH = 7.2) or FBS (500 μL) at 37°C for 2 h while shaking at 400 rpm. The reaction progression was monitored by radio-TLC (iTLC) with 0.1 M citrate buffer (pH 5.0) as the mobile phase. Under these conditions, radioimmunoconjugates remained stationary at the origin, while [^89^Zr]Zr-DFO displayed an R_f_ of 0.2-0.3.

### CD69 and CD20 plasmid design and lentivirus production

The murine CD69 sequence was acquired from Origine (MR226790L4) and cloned into a pHIV.EGFP vector to create pHIV.CD69.IRES.EGFP. The CD20 DNA sequence was acquired from the Gill Lab at the University of Pennsylvania and synthesized by IDT. The CD20 insert was then cloned into the pTRPE vector to create pTRPE.CD20.

For the lentiviral production of the pHIV.CD69.IRES.EGFP vector, VSV-g (7 µg), pRSV-Rev (18 µg), pGag/Pol (18 µg), and 15 µg of vector of interest (pHIV.CD69.IRES.EGFP) were used. For virus production, HEK293T/17 cells were plated a day prior to transfection with the packaging plasmids and vector of interest mentioned above in a T-150 flask. On the day of transfection, a half-media change was conducted and the Lipofectamine 3000 protocol for transfection was followed using the materials mentioned along with 90 µL of Lipofectamine 3000 and 30 µL P3000 (Invitrogen, L3000001). Supernatants were collected 24, 48, and 72 h post transfection. Supernatants were then centrifuged to remove cell debris and filtered through a 0.45 µm PVDF membrane (Millipore Sigma, SE1M003M00) prior to concentration using a 100 kDa Amicon centrifugal filter concentrator (Millipore Sigma, UFC910096). Concentrated lentiviruses for each construct were stored at −80°C.

For the lentiviral production of the pTRPE.CD20 vector, pRSV/Rev (18 µg), pGag/Pol (18 µg), and pCocal-g (3 µg), and 27 µg of vector of interest (pTRPE.CD20) were used. For virus production, HEK293T/17 cells were plated a day prior to transfection with packaging plasmids in a T-150 flask. On the day of transfection, a half-media change was conducted and the Lipofectamine 2000 protocol for transfection was followed using the materials mentioned along with 90 µL of Lipofectamine 2000 (Invitrogen, 11668019). Virus was prepared and stored as detailed previously.

### Cell line generation and culture

The Raji and CT26 cells utilized in this study were obtained from ATCC (CCL-86 and CRL-2638). Culturing was performed in accordance with ATCC guidelines, using RPMI-1640 Medium supplemented with 2 mmol/L L-glutamine, 10 mmol/L HEPES, 1 mmol/L sodium pyruvate, 4,500 mg/L glucose, and 1,500 mg/L sodium bicarbonate (Gibco, A1049101). The culture medium was further enriched with 10% FBS (Gibco, 16000044) and 1% penicillin/streptomycin (Invitrogen, 15140122). The cells were maintained in a humidified incubator at 37°C. HEK293T/17 cells (ATCC, CRL-11268) were cultured in complete media: DMEM containing 10% fetal bovine serum (Invitrogen), 2 mM glutamine (Gibco), 1% penicillin/streptomycin (Gibco), and were maintained in a humidified incubator at 37°C. To ensure optimal performance, cells were utilized for *in vitro* and *in vivo* assays within 3 months of thawing, and passaging was limited to approximately 10 times prior to experimentation.

To generate CT26-CD69 cells, pHIV.CD69.IRES.EGFP lentivirus and 10 µg/mL polybrene was added to CT26 cells in antibiotic-free complete culture medium. Cells were then sorted by the Penn Cytomics & Cell Sorting Shared Resource Laboratory at the University of Pennsylvania (RRid:SCR_022376). To generate I45-CD20 cells, pTRPE.CD20 lentivirus and 10 µg/mL polybrene was added to I45 cells in antibiotic free complete culture medium, and expression was confirmed using an AF647-conjugated anti-CD20 antibody and an LSR II cytometer (BD Biosciences) in the Penn Cytomics & Cell Sorting Shared Resource Laboratory.

### Rituximab-AF647 conjugation

Alexa Fluor 647 (AF647) was conjugated to rituximab (∼100 mg) using an antibody labeling kit (Life Technologies, A20186). AF647-conjugated rituximab antibodies were used for flow cytometry assays to validate CD20 expression on I45-CD20 cells.

### In vitro demonstration of TKO strategy

Raji (CD20^+^) and K562 cells (CD20^-^) in suspension (5 x10^6^ cells in 1 mL 2% BSA/Opti-MEM) were incubated with [^89^Zr]Zr-DFO-TCO-rituximab at 37°C for 2 h and subsequently washed three times with 1% DMSO in PBS. To assess nonspecific binding, excess unlabeled rituximab (100 μg/mL) was introduced simultaneously with the radiotracer. Cell pellets were then collected by centrifugation and radioactive uptake was measured using a gamma counter (Wizard Detector, Perkin Elmer). Uptake (%ID/million cells) was calculated by dividing measured radioactivity by the starting incubated dose of [^89^Zr]Zr-DFO-TCO-rituximab and normalizing to sample cell counts. Following a similar experimental protocol, CT26-CD69^+^ and CT26-WT cells (5 x 10^6^) were incubated with [^89^Zr]Zr-DFO-TCO-H1.2F3 at 25°C for 1 h in 1 mL Opti-MEM supplemented with 2% BSA. This was done in the presence and absence of the parent H1.2F3 antibody (100 μg/mL).^33^ Uptake was assessed using the previously described method. For the Tz cleavage study, Raji cells (CD20^+^) and CT26-CD69^+^ cells were further treated with Tz-1 (1 mg/mL) in 1% DMSO/Opti-MEM for 1 h. The cells were then thoroughly washed with 1% DMSO/PBS, centrifuged to collect cell pellets, and counted using a gamma counter.

### Healthy mouse biodistribution

[^125^I]I-AG-TCO-rituximab (0.93 MBq) was administered through the tail vein in healthy female Balb/c mice (6 weeks old, n = 6). After a 3 h period, the mice were randomly assigned to two groups. One group was treated with Tz-1 (1 mg in 5% EtOH-PBS/mice), while the other received a vehicle (5% EtOH/PBS). Following this, the mice were anesthetized with 2% isoflurane and euthanized for biodistribution studies 1 h after Tz or vehicle administration. Various organs of interest and blood samples were collected, and the uptake in each organ was evaluated using a gamma counter (PerkinElmer). The results were quantified as a percentage of injected dose per gram of organ (%ID/g). For urine analysis, samples were collected immediately after euthanizing the animals and subjected to radio-HPLC (Agilent 1260 Infinity II HPLC with Agilent column: 300SB-C18, 9.4 X 250 mm, 5 µm). The mobile phase started with 95% solvent A (0.1% TFA in water) and 5% solvent B (0.1% TFA in acetonitrile), transitioning to 50% solvent A at 10 minutes, maintaining that condition for 2 minutes, and finally returning to the initial condition in 15 minutes. The flow rate was maintained at 1 mL/min during the analysis.

### Small animal PET/CT imaging

Experimentation involving tumor models and small animal PET imaging was conducted in accordance with ethical principles and approved protocols outlined by the Institutional Animal Care and Use Committee (IACUC) Office of Animal Welfare at the University of Pennsylvania (IACUC protocol #805447). Female CD1 nu/nu mice, aged six to eight weeks, were subcutaneously xenografted with 1 x 10^6^ Raji or I45-CD20^+^ cells, suspended in 75 μL RPMI complete medium and 75 μL Matrigel Matrix (Corning, 354234), on the right flank. After three weeks of tumor growth, reaching an approximate volume of 100-120 mm^3^, animals were intravenously administered [^89^Zr]Zr-DFO-TCO-rituximab (7.4-9.3 MBq) or [^68^Ga]Ga-NOTA-TCO-rituximab (18.5-22 MBq). Post-antibody administration, animals were randomly allocated into two groups. One group received Tz-1 (1 mg in 5% EtOH-PBS/mouse), while the other was administered a vehicle (5% EtOH/PBS). Subsequently, animals were anesthetized with 2% isoflurane for PET/CT imaging on a small animal PET/CT system (Molecubes) 1 h after Tz or vehicle administration. PET/CT image analysis was executed using MIM Software (Version 7.1.5). In brief, regions of interest (ROI) were demarcated on tumors and selected background organs for the determination of SUV_max_ and SUV_mean_ values. Tumor/background ratios were then calculated by dividing tumor SUV_max_ by organ SUV_max_ or tumor SUV_mean_ by organ SUV_mean_.

### Ex vivo biodistribution

The mice were euthanized following [^89^Zr]Zr-DFO-TCO-rituximab or [^68^Ga]Ga-NOTA-TCO-rituximab PET/CT imaging. Tissues were collected, and the uptake in each organ was assessed using a gamma counter (PerkinElmer), quantifying the results as a percentage of injected dose per gram of organ (%ID/g).

## Author Contributions

S.S. and M.A.S. conceived the project. S.S., N.S. A.P.K and A.F.D.C. performed chemical synthesis and characterization of the molecules. S.S., J.P., K.J.E., N.S., K.X. and A.P.K. performed *in vitro* experiments, analyzed data, and interpreted results. S.S., J.P., K.J.E., K.X. performed *in vivo* animal experiments and interpreted the results with M.A.S. S.S. and J.P. wrote the first draft manuscript. All other authors contributed to the final draft.

## Acknowledgments

We would like to thank the Small Animal Imaging Facility (Eric Blankemeyer), the University of Pennsylvania Cyclotron and Radiochemistry Facility (Alexander Schmitz and Sharon Lee), and the Penn Cytomics and Cell Sorting Shared Resource Laboratory (RRID: SCR_022376) for making this research possible. We thank the University of Pennsylvania Chemistry Department for NMR and LCMS support, Mayne Leland for recording HRMS, as well as David Mankoff and Robert H. Mach for helpful discussions.

## Abbreviation

AG: 4-amino-2-iodophenol
NOTA: *2,2′,2”-(1,4,7-triazacyclononane-1,4,7-triyl)triacetic acid*
DFO: Deferoxamine
ID: Injected dose
SUV: standardized uptake values
EDTA: Ethylenediaminetetraacetic acid.

## Additional information

Supplementary information The online version contains supplementary material available at

