## Supplemental File for "A general approach to reduce off-target radioactivity in vivo via Tetrazine-Knock-Out (TKO)"

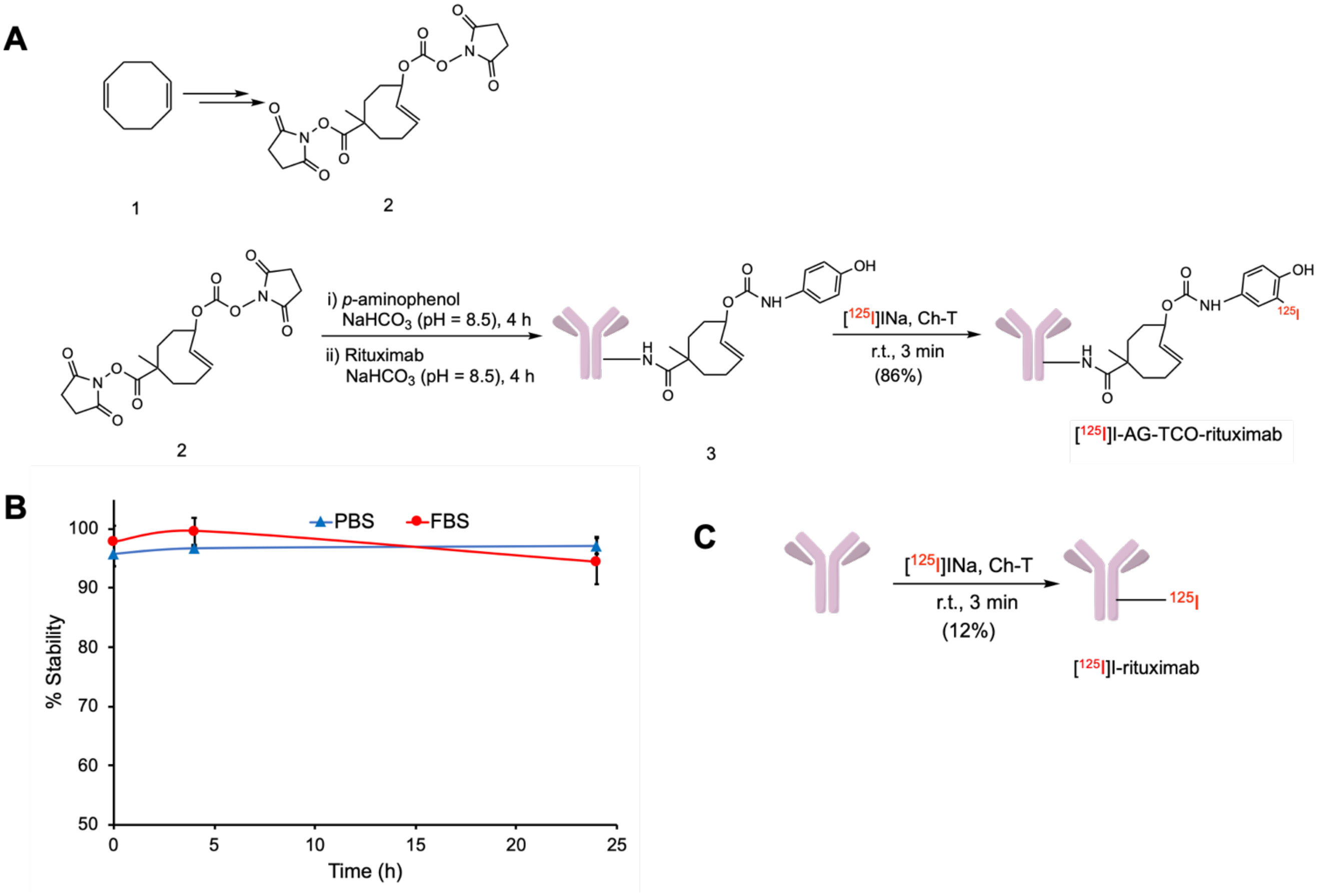


**Fig S1. A**) Synthesis of [$^{125}$I]I-AG-TCO-rituximab. **B**) Serum stability of [$^{125}$I]I-AG-TCO-rituximab. Radioimmunoconjugate (3.7 MBq) was incubated with PBS (pH = 7.4) or FBS (500 μL) at 37 °C for 24 h and stability was measured by radio-TLC (iTLC/saline). In the iTLC-saline system, free I-125 moved to the solvent front ($R_f$ = 1), while radio-immunoconjugate remained at the origin ($R_f$ = 0); **C**) Radioiodination of the parent rituximab under same radiolabeling condition.

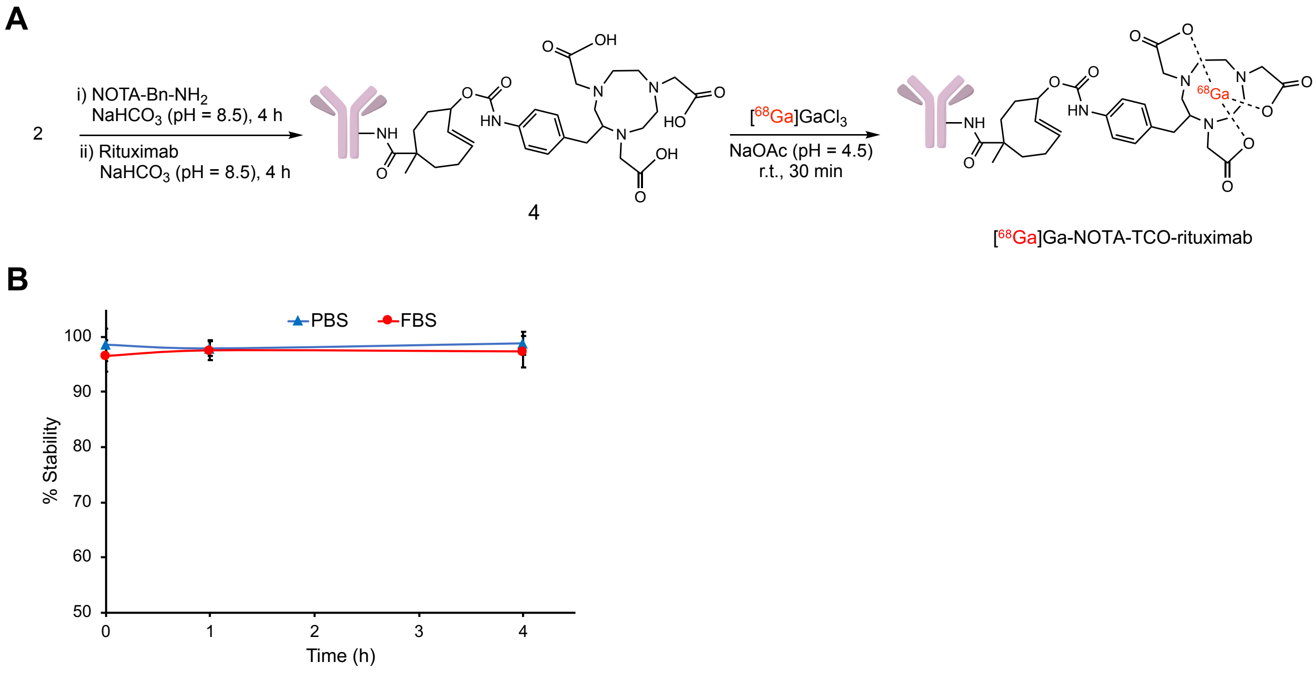


**Fig S2. A**) Synthesis of [$^{68}$Ga]Ga-NOTA-TCO-rituximab. **B**) Serum stability of [$^{68}$Ga]Ga-NOTA-TCO-rituximab. Radioimmunoconjugate (3.7 MBq) was incubated with PBS (pH = 7.4) or FBS (500 μL) at 37°C for 24 h and stability was measured by radio-TLC (iTLC-SA/50 mM EDTA (pH 4.5). In the ITLC-SA system, free Ga-68 formed a complex with EDTA and eluted with the solvent front ($R_f$ = 1), while radio-immunoconjugate remained at the origin ($R_f$ = 0).

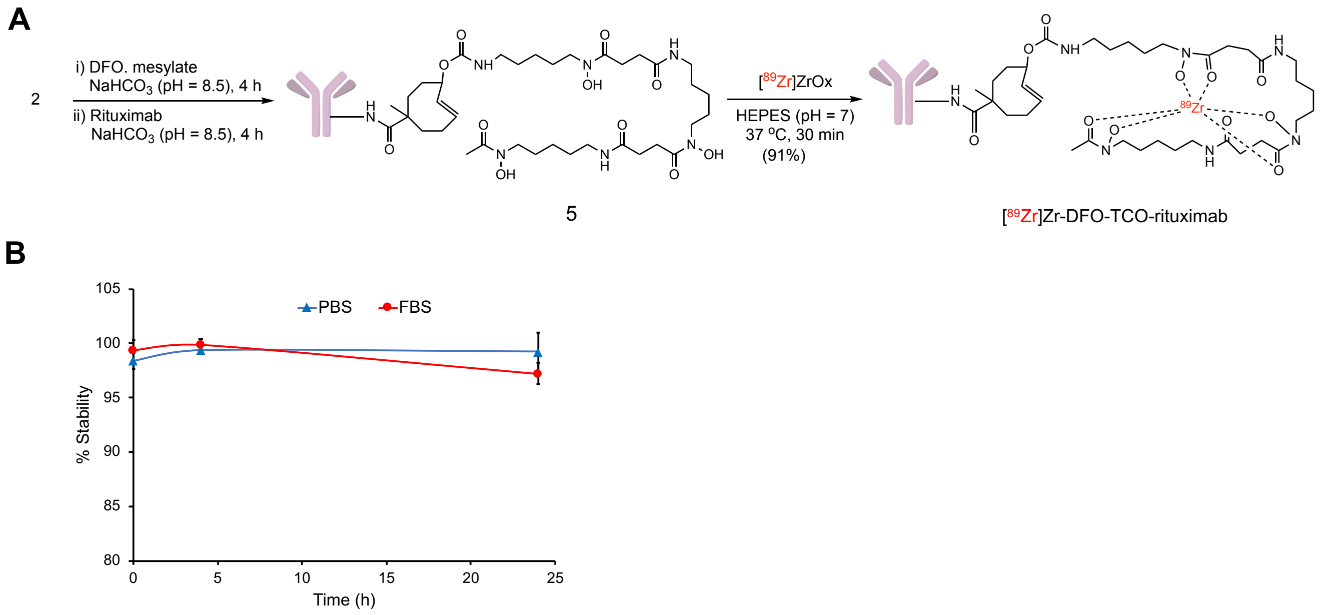


**Fig S3. A**) Synthesis of [$^{89}$Zr]Zr-DFO-TCO-rituximab. **B**) Serum stability of [$^{89}$Zr]Zr-DFO-TCO-rituximab. Radioimmunoconjugate (3.7 MBq) was incubated with PBS (pH = 7.4) or FBS (500 mL) at 37°C for 24 h and stability was measured by radio-TLC (iTLC-SA/0.1 M EDTA (pH 5). In the ITLC-SA system, free Zr-89 formed a complex with EDTA and eluted with the solvent front ($R_f$ = 1), while radio-immunoconjugate remained at the origin ($R_f$ = 0).

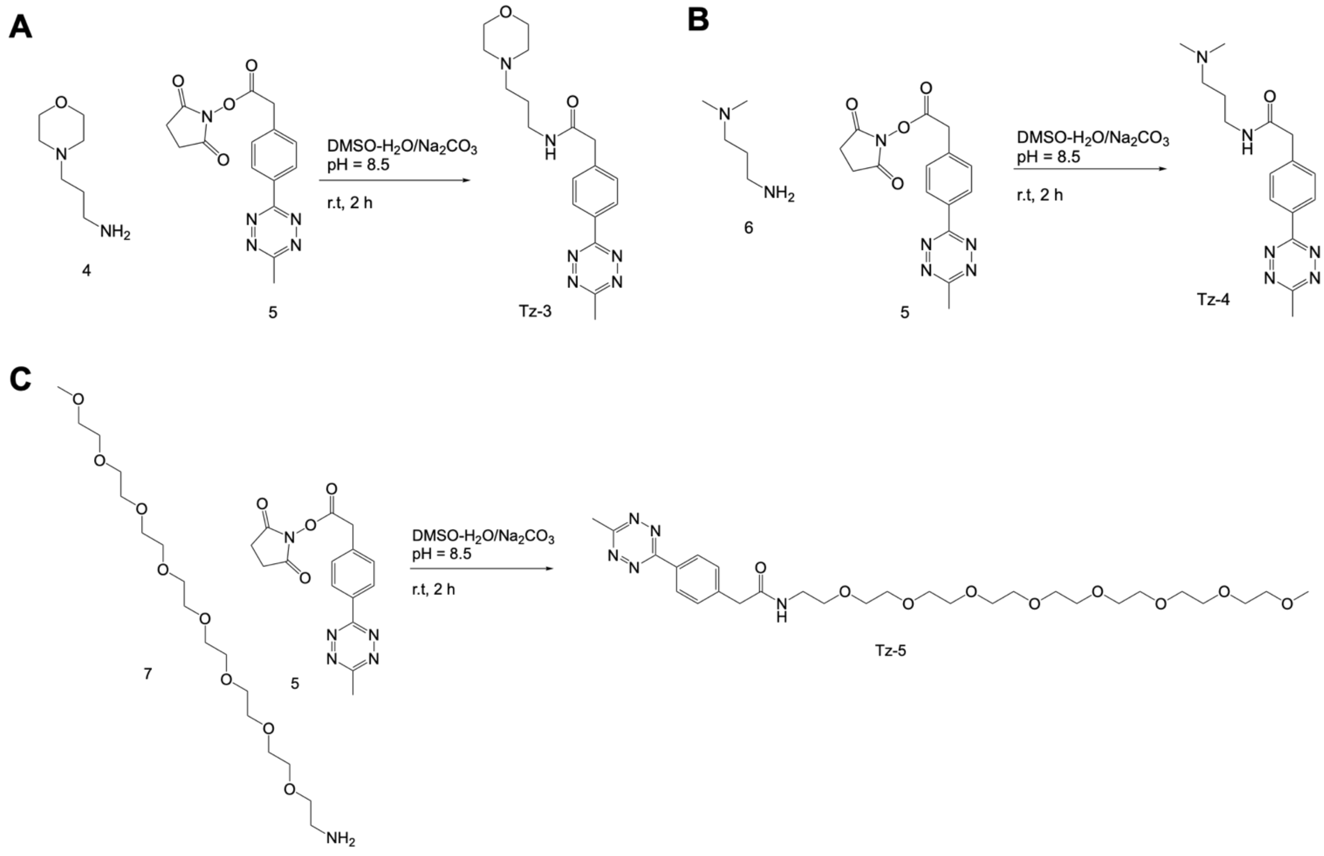


**Fig S4.** Synthesis of **A**) Tz-3, **B**) Tz-4 and **C**) Tz-5

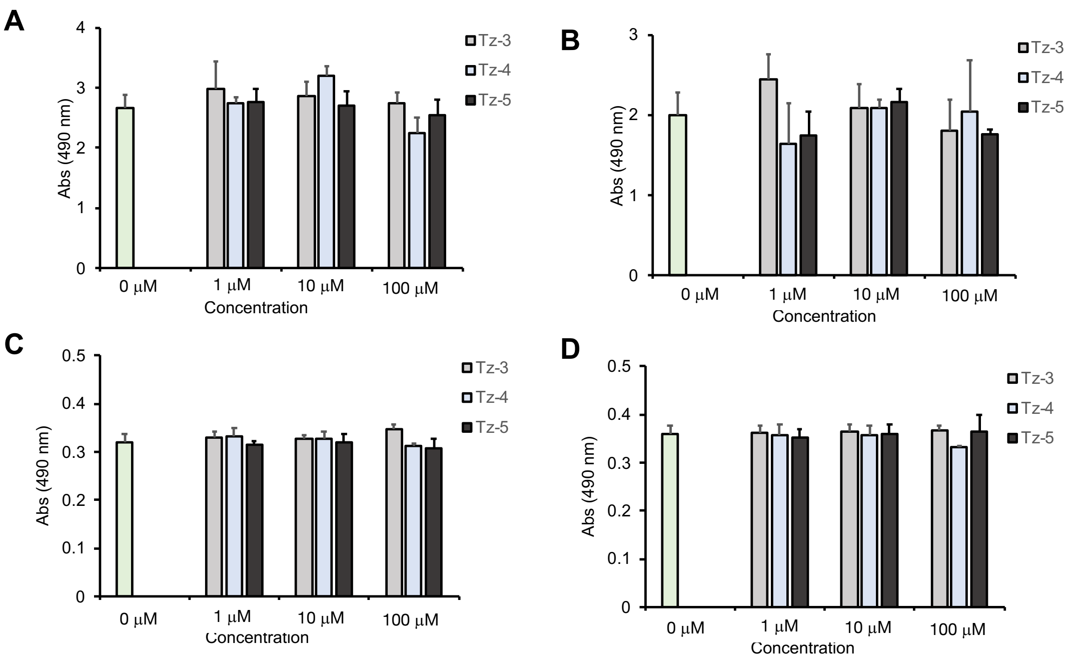


**Fig S5.** Cell viability assays for Tz-3-5. HEK293T/17 (**A & B**) and primary hepatocyte (**C & D**) cells were incubated with Tz-3-5 for 1 h (**A & C**) or 4 h (**B & D**) at 37 °C in 5% $CO_2$. 1% DMSO in media supplemented with 5% FBS and 1% penicillin/streptomycin was used as a control. Briefly, 4E5 cells were plated in 96 well plates in complete media. After 24 h, cells were treated with Tz-3-5 or vehicle for an additional 1-4 h. Cell viability was subsequently assessed using the Promega CellTiter 96® AQueous One Solution Cell Proliferation Assay and absorbance at 490 nm was recorded using a microplate reader (ThermoFisher Varioskan).

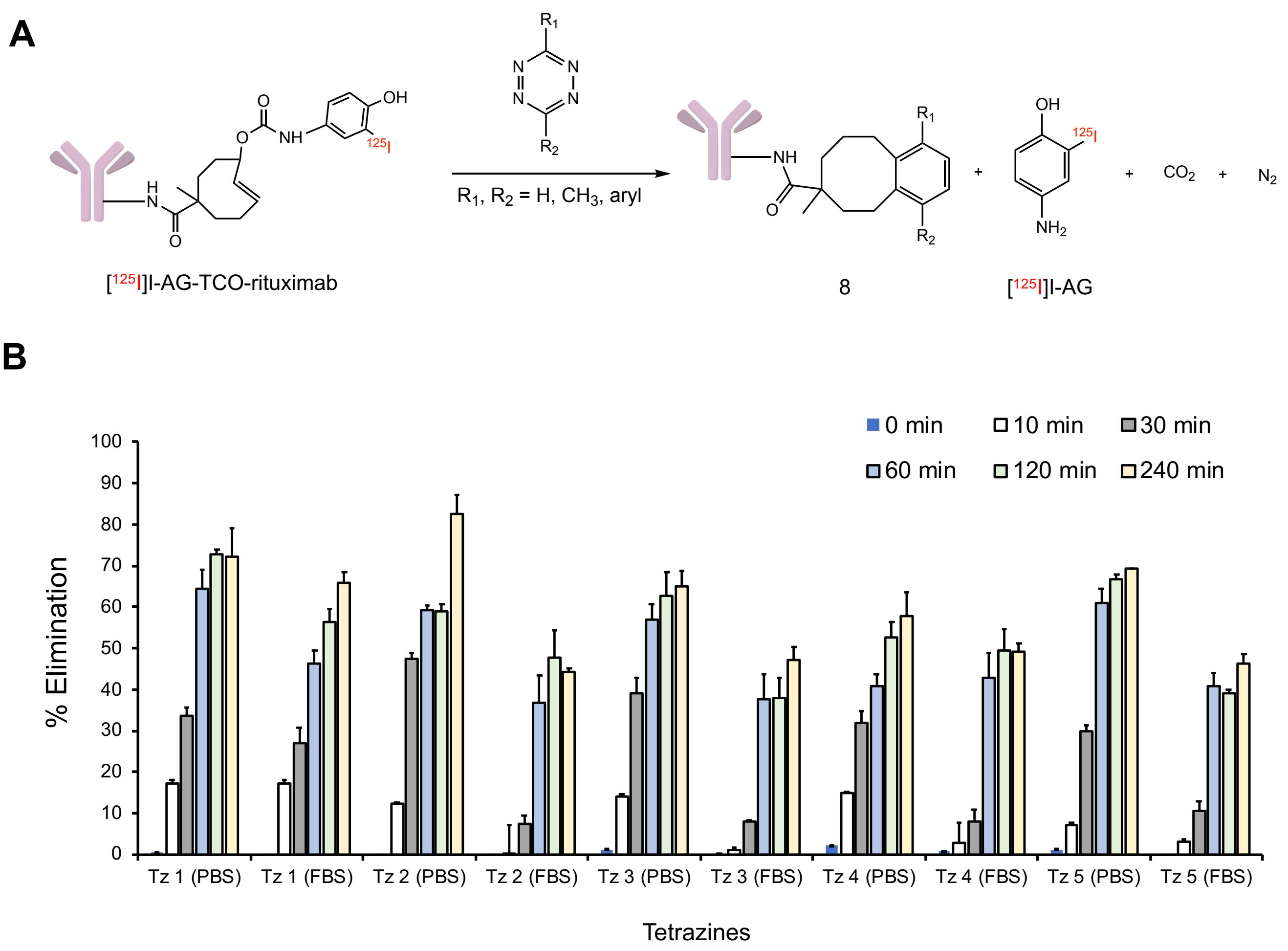


**Fig S6.** *In vitro* knock out studies. **A**) [$^{125}$I]I-AG release from [$^{125}$I]I-AG-TCO-rituximab upon reaction with tetrazines **Tz-1-5**. **B**) Release of [$^{125}$I]I-AG from [$^{125}$I]I-AG-TCO-rituximab in PBS (pH = 7.4) and FBS (500 μL) at 37°C up to 4 h. The progress of the reaction was monitored by radio-TLC (silica, 5% MeOH/DCM). The data represent the mean ± SD (n = 3).

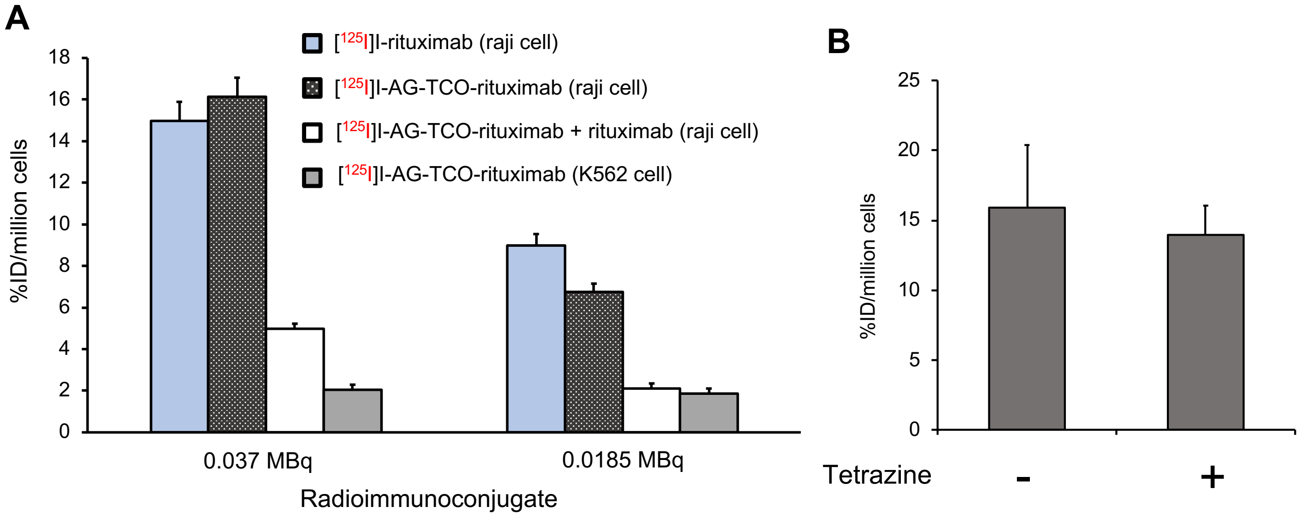


**Fig S7.** Demonstration of the TKO strategy in cells with rituximab. **A**) Cell uptake study of [$^{125}$I]I-rituximab and [$^{125}$I]I-AG-TCO-rituximab in Raji (CD20$^{+}$) and K562 cells (CD20$^{-}$). Cells were incubated with the radioimmunoconjugates at 37°C for 2 h in Opti-MEM supplemented with 2% BSA in the presence or absence of parent rituximab (100 μg). Uptake was counted in a γ–counter (n = 3); B) TKO strategy with an internalizing antibody. Raji cells were first incubated with [$^{125}$I]I-AG-TCO-rituximab in Opti-MEM supplemented with 2% BSA and then incubated in presence and absence of Tz-1 for 1 h for selective release of [$^{125}$I]I-AG. Cells were then thoroughly washed with 1% DMSO/PBS, centrifuged and counted in a γ-counter (n = 3). No noticeable difference was observed in antibody retention in both Tz-treated and untreated cells.

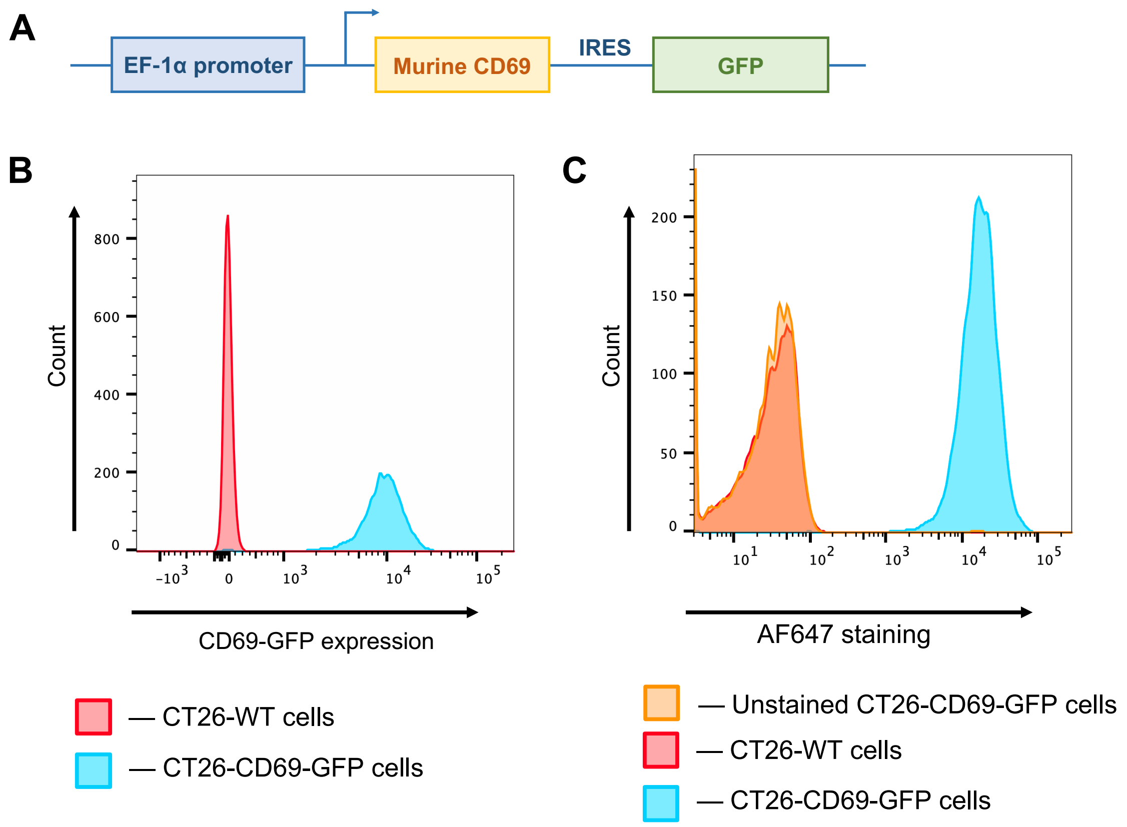


**Fig S8.** Characterization of CT26-CD69 cells. **A**) Scheme showing pHIV-CD69-GFP vector; **B**) Flow cytometry analysis showing CD69 expression in CT26 cells that were transduced to express CD69-GFP relative to the CT26 WT negative control, **C**) Flow cytometry analysis of CT26-CD69 and CD26-WT cells that have been stained with an AF647-conjugated anti-mCD69 full length antibody (AF647-H1.2F3).

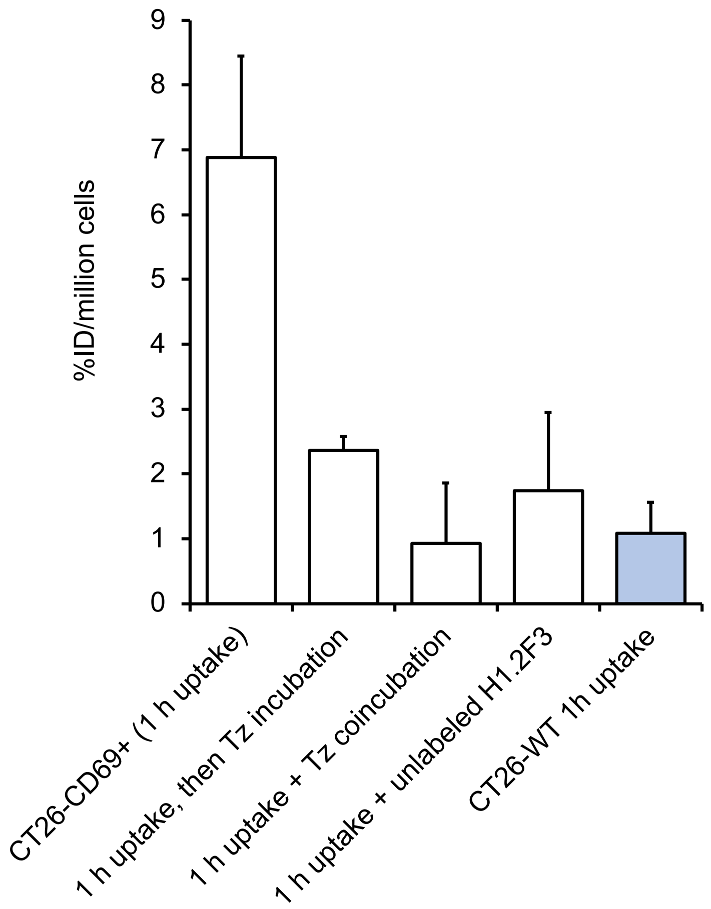


**Fig S9.** Demonstration of TKO with a slowly internalizing antibody. CT26-CD69$^+$ and CT26-WT cells were incubated with [$^{125}$I]I-AG-TCO-H1.2F3 at 25°C for 1 h in Opti-MEM supplemented with 2% BSA in presence and absence of parent H1.2F3 antibody (100 μg). CT26-CD69$^+$ cells were then incubated with Tz-1 for 1 h for selective release of [$^{125}$I]I-AG followed by thorough washing with 1% DMSO/PBS, centrifugation to collect the cell pellets, and counted in a γ -counter (n = 3). Tz coincubation was done with [$^{125}$I]I-AG-TCO-H1.2F3 in CT26-CD69$^+$ cells under the same condition. Cells were thoroughly washed with 1% DMSO/PBS as earlier before counted in a γ-counter (n = 3). A significant drop in cellular uptake was noticed in Tz-treated cells, confirming the release of the radioactive counterpart from the antibody.

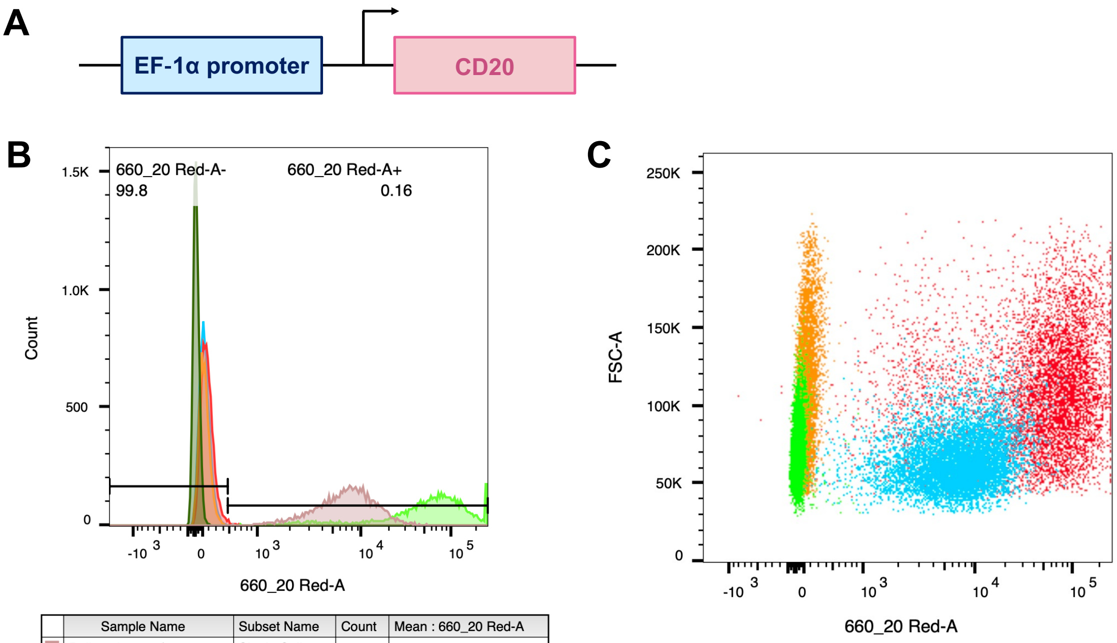


| | Sample Name | Subset Name | Count | Mean : 660_20 Red-A |
|---|---|---|---|---|
| | Raji_stain_002.fcs | Single Cells | 7939 | 8899 |
| | Raji_US_001.fcs | Single Cells | 7694 | 37.5 |
| | I45_CD20 stain_006.fcs | Single Cells | 7110 | 73334 |
| | I45_CD20 US_005.fcs | Single Cells | 6200 | 151 |
| | I45_WT stain_004.fcs | Single Cells | 8013 | 223 |
| | I45_WT US_003.fcs | Single Cells | 7335 | 155 |

| | Sample Name | Subset Name | Count |
|---|---|---|---|
| | Raji_US_001.fcs | Single Cells | 7694 |
| | Raji_stain_002.fcs | Single Cells | 7939 |
| | I45_CD20 US_005.fcs | Single Cells | 6200 |
| | I45_CD20 stain_006.fcs | Single Cells | 7110 |

**Fig S10. A**) Scheme showing CD20 vector; **B** and **C**) Flow cytometry analysis verifying the expression of the CD20 protein on the surface of Raji and I45-CD20 cells. Expression of CD20 on the cells was assessed after staining with AF647-conjugated rituximab. Unstained (US) and I45-WT cells served as negative controls.

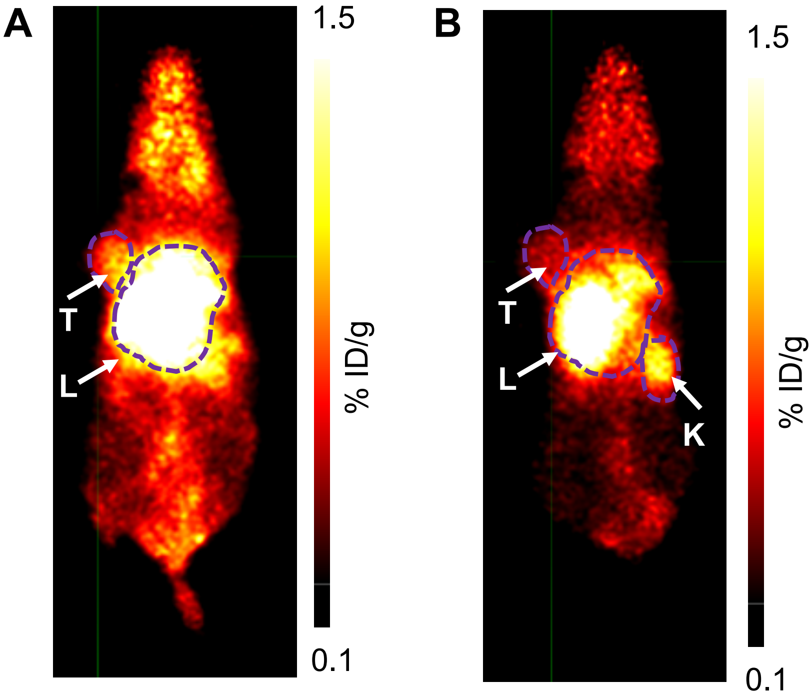


**Fig S11.** Female nu/nu mice bearing I45-CD20$^{+}$ xenografts were administered with [$^{68}$Ga]Ga-NOTA-TCO-rituximab and imaging and biodistribution studies were conducted as described in **Fig 5**. **A**) Representative PET image (coronal) for the vehicle-treated group (n = 3); **B**) Representative PET image (coronal) for the Tz treated group (n = 3).

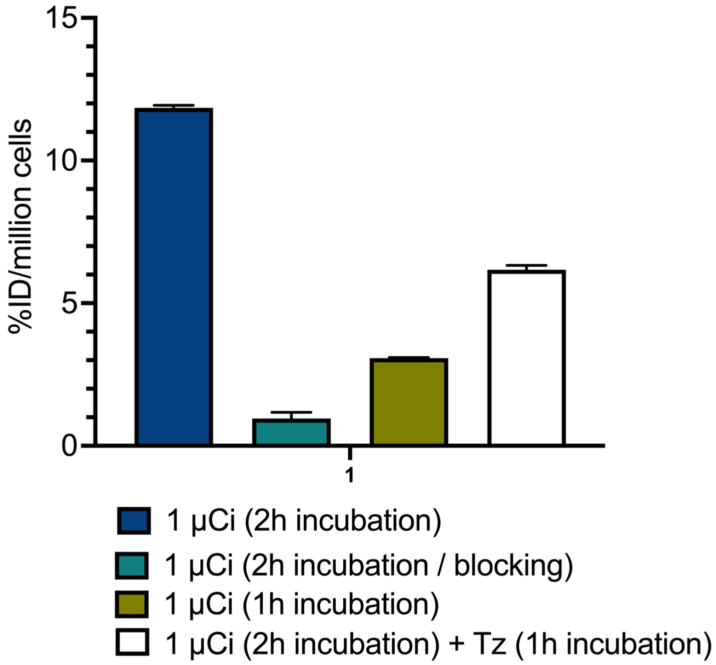


**Fig S12.** Demonstration of the TKO strategy in cells in I45-CD20$^+$ cells. Cell uptake study of [$^{68}$Ga]Ga-NOTA-TCO-rituximab was performed by incubating the cells with the radioimmunoconjugates at 37°C in Opti-MEM supplemented with 2% BSA in presence or absence of parent rituximab (100 µg). Uptake was counted in a γ-counter (n = 3). Cells were then incubated with Tz-1 for 1 h for selective release of [$^{68}$Ga]Ga-NOTA. Cells were then thoroughly washed with 1% DMSO/PBS, centrifuged and counted in a γ-counter (n = 3).

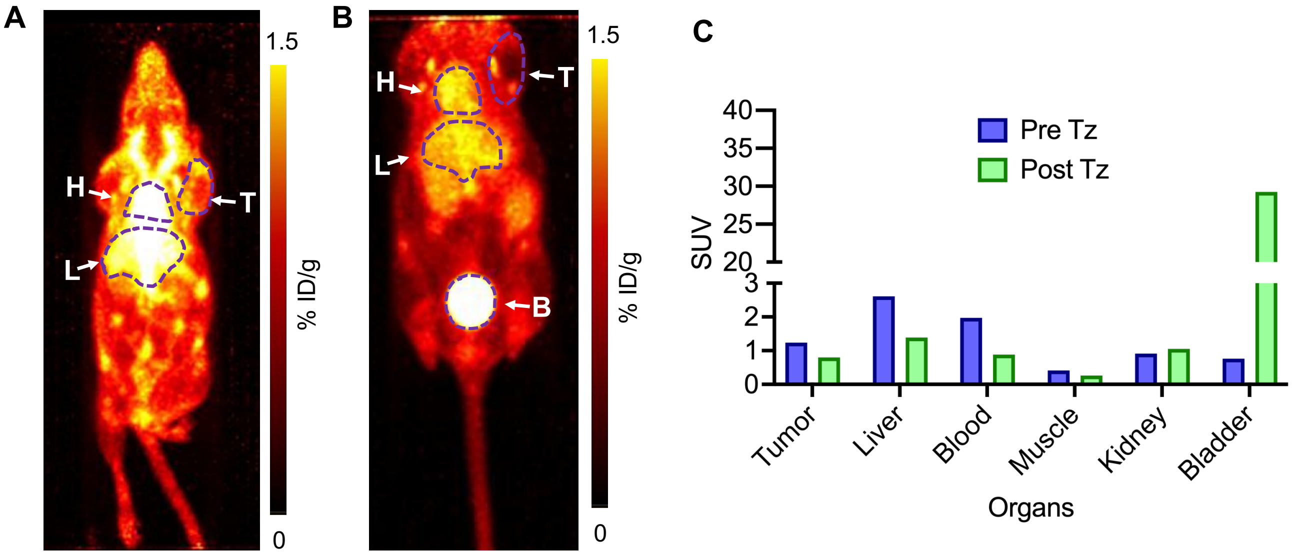


**Fig S13.** A female nu/nu mouse bearing with Raji xenografts was injected with [$^{89}$Zr]Zr-DFO-TCO-obinutuzumab and subsequently underwent microPET scans 24 h post-injection. Following this imaging, the same mouse was given Tz and scanned again using microPET 1 h after Tz administration. **A**) MIP images before Tz administration and **B**) after Tz administration. **C**) SUV for various tissues including tumor, liver, blood, muscle, kidney, and bladder were calculated before and after Tz administration in the same animal.

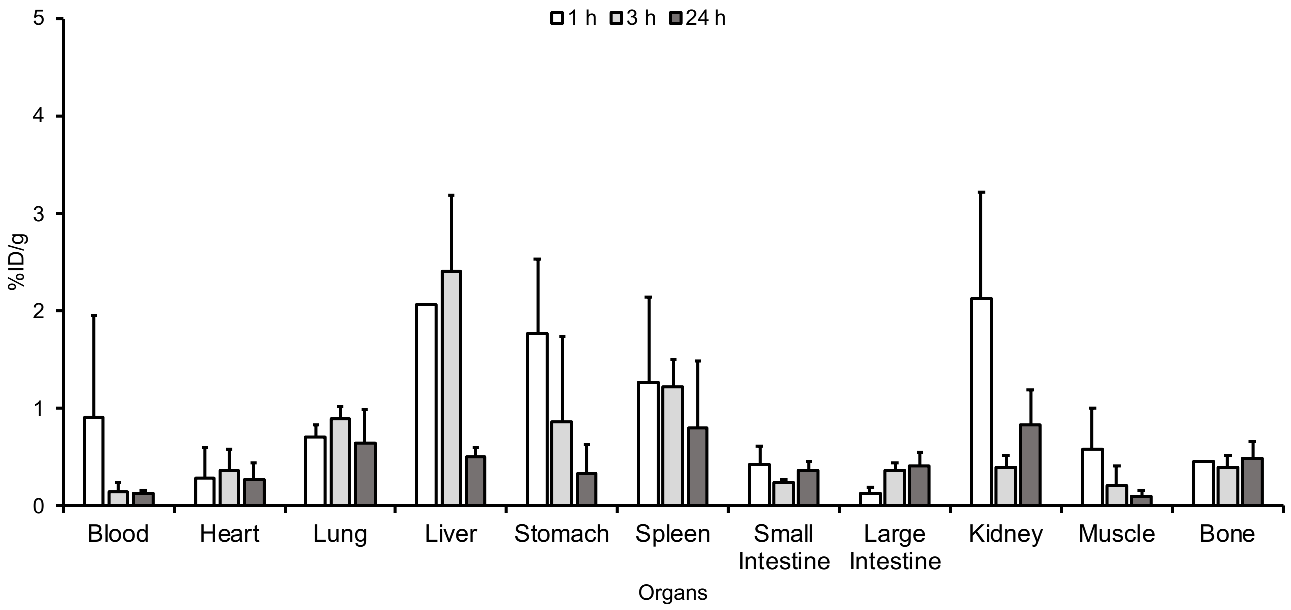


**Fig S14.** Bio-distribution of [$^{89}$Zr]Zr-DFO at 1, 3 and 24 h post-injection in healthy female Balb/c mice (6 weeks, n = 3 for each group). [$^{89}$Zr]Zr-DFO (0.74 MBq, 100μL) was injected via the tail vein and animals were sacrificed under anesthesia (isoflurane). Blood and various tissues of interest were collected (cotton gauze was used to remove excess blood), weighed, and counted in a γ-counter (Perkin Elmer). To determine the percentage of the injected dose, the samples were compared with a diluted standard solution obtained from the injected solution. The results were expressed as the percentage of the injected dose per gram (%ID/g) of wet tissue.

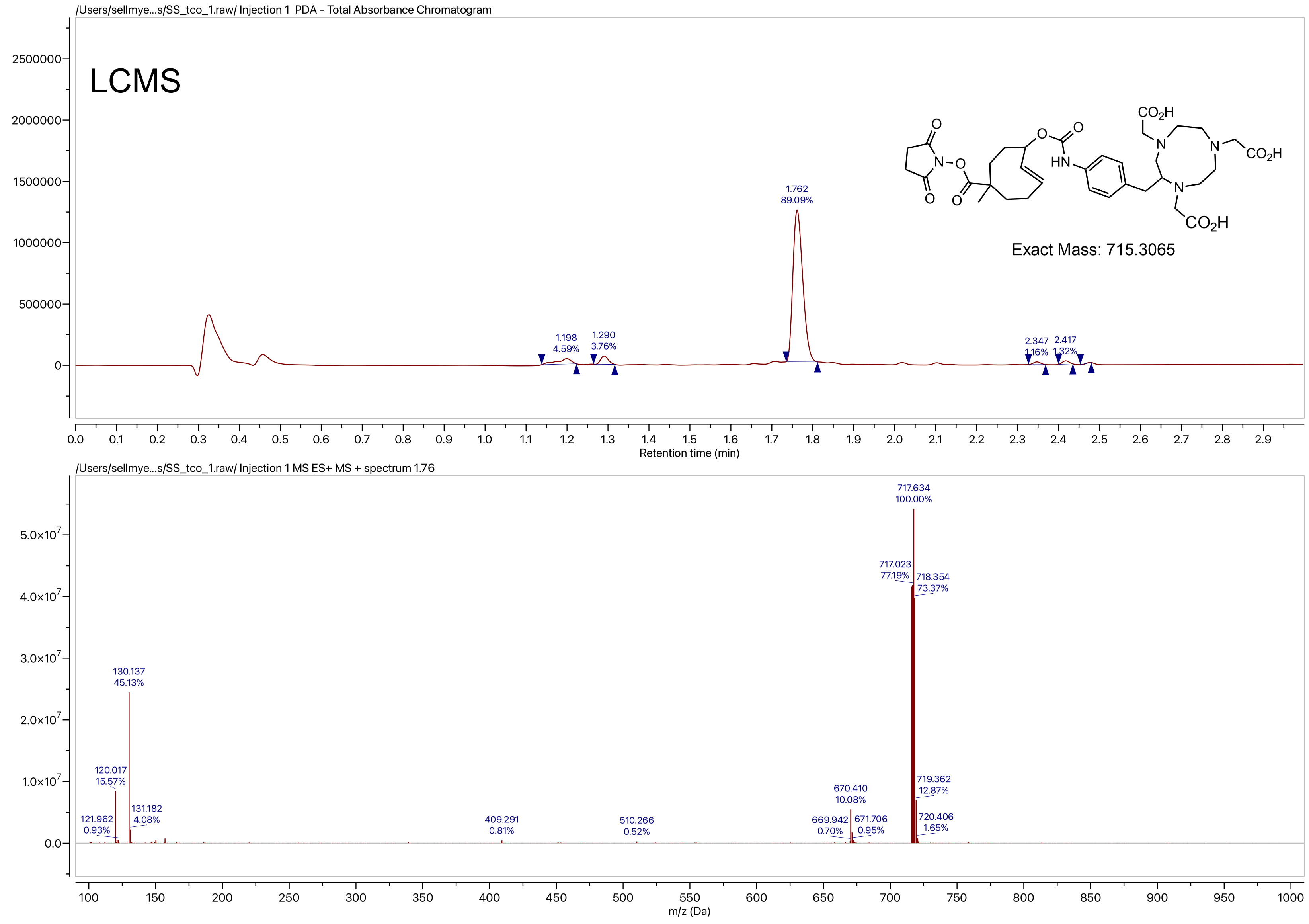
/Users/sellmye...s/SS_tco_1.raw/ Injection 1 PDA - Total Absorbance Chromatogram
LCMS
Exact Mass: 715.3065
1.762
89.09%
1.198
4.59%
1.290
3.76%
2.347
1.16%
2.417
1.32%
Retention time (min)
/Users/sellmye...s/SS_tco_1.raw/ Injection 1 MS ES+ MS + spectrum 1.76
717.634
100.00%
717.023
77.19%
718.354
73.37%
719.362
12.87%
720.406
1.65%
670.410
10.08%
669.942
0.70%
671.706
0.95%
510.266
0.52%
409.291
0.81%
130.137
45.13%
120.017
15.57%
121.962
0.93%
131.182
4.08%
m/z (Da)

SS_Tz_Mor.1.fid

### $^{1}H$ NMR, 400 MHz, DMSO-$d_6$

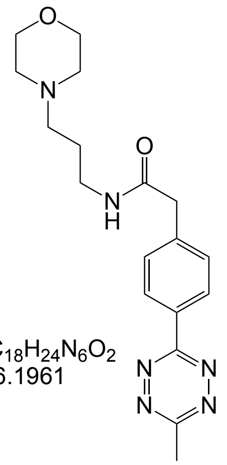


Chemical Formula: $C_{18}H_{24}N_6O_2$
Exact Mass: 356.1961

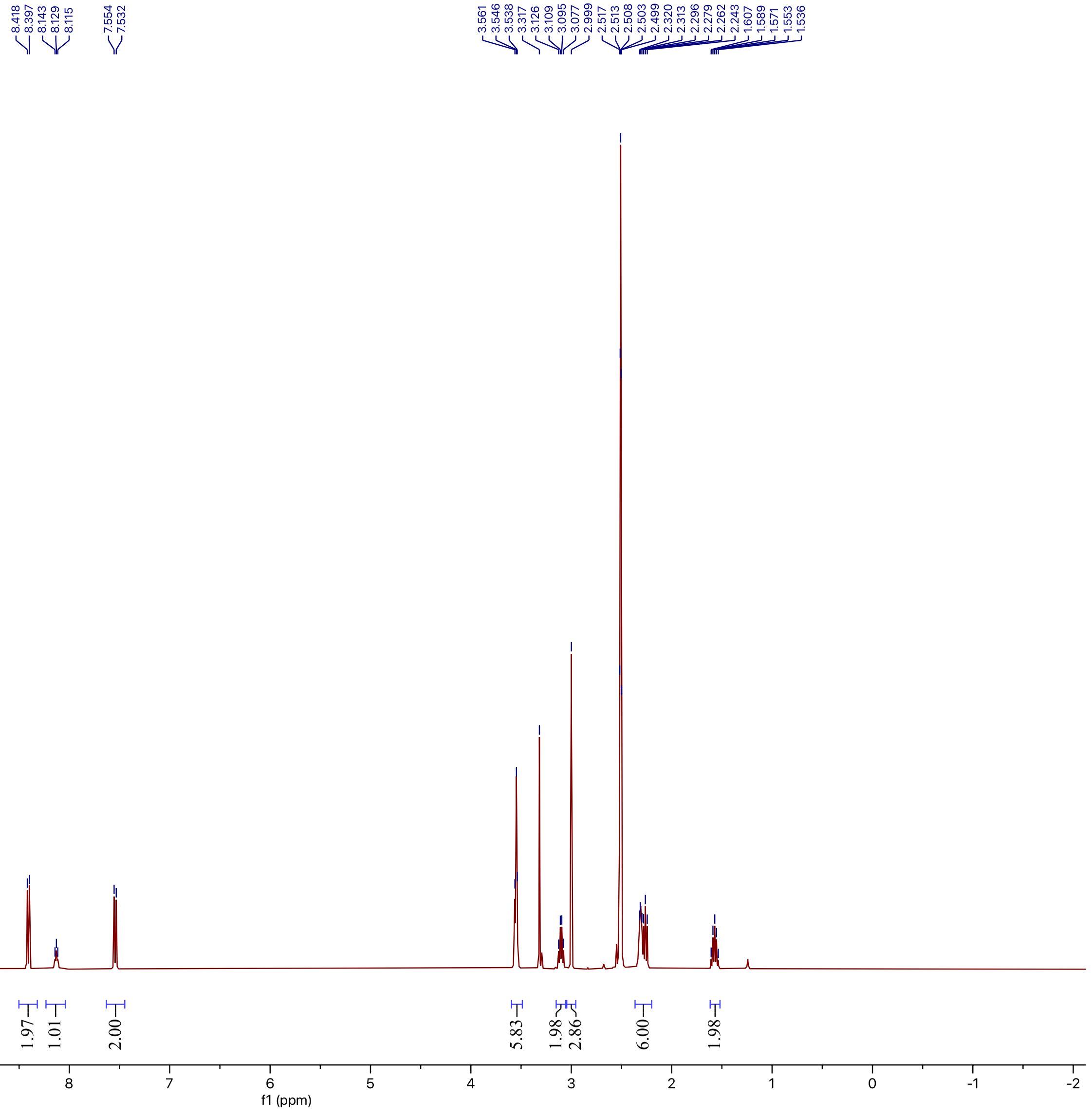

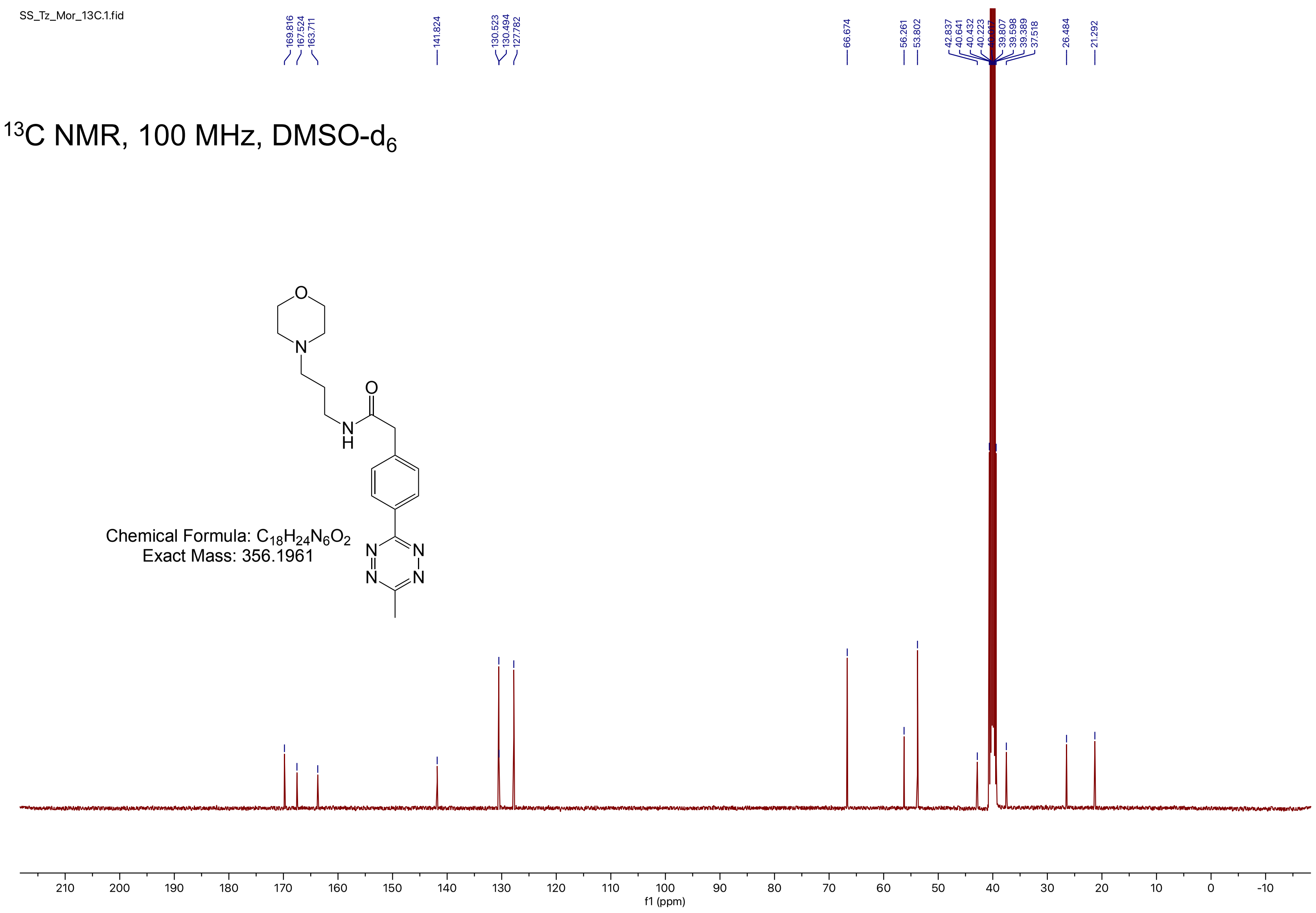
SS_Tz_Mor_13C.1.fid
169.816
167.524
163.711
141.824
130.523
130.494
127.782
66.674
56.261
53.802
42.837
40.641
40.432
40.223
39.807
39.598
39.389
37.518
26.484
21.292
13C NMR, 100 MHz, DMSO-d6
Chemical Formula: C18H24N6O2
Exact Mass: 356.1961
O
N
H
f1 (ppm)

HRMS

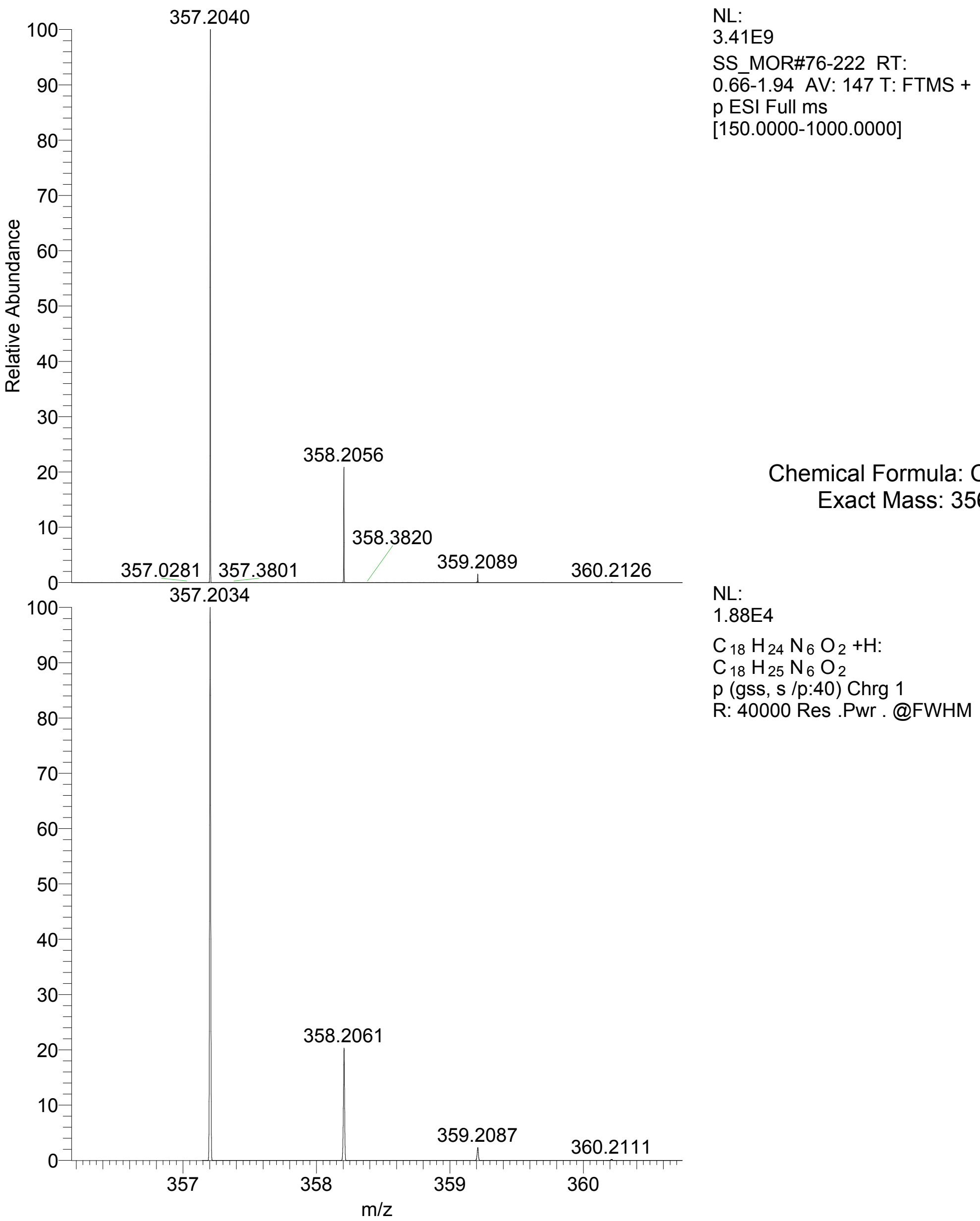


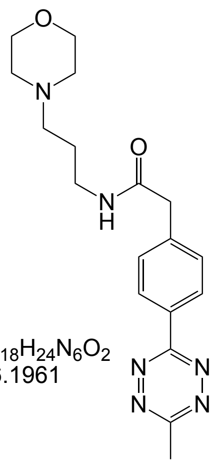

Chemical Formula: $C_{18}H_{24}N_6O_2$
Exact Mass: 356.1961

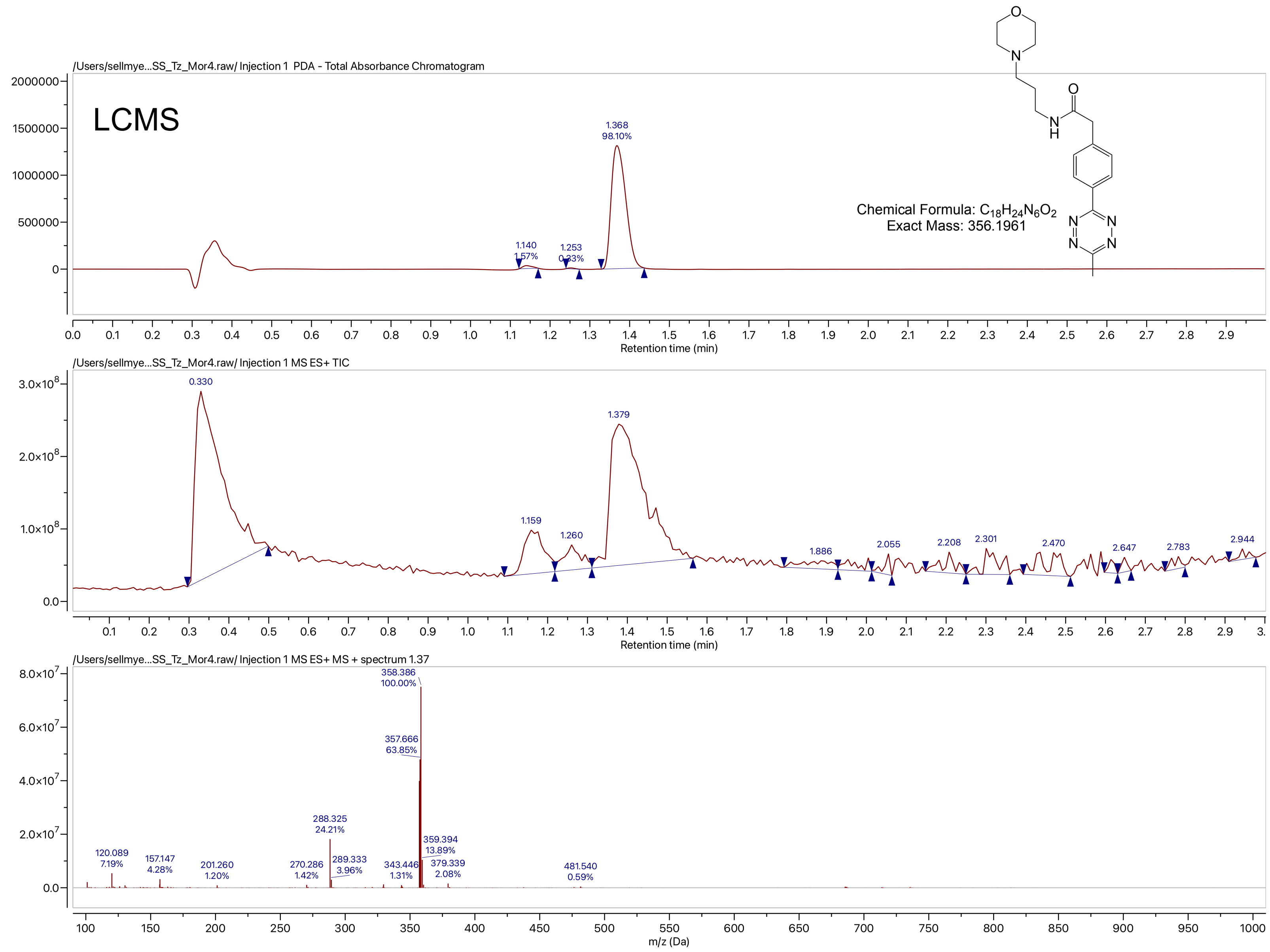
LCMS
Chemical Formula: $C_{18}H_{24}N_6O_2$
Exact Mass: 356.1961
/Users/sellmye...SS_Tz_Mor4.raw/ Injection 1 PDA - Total Absorbance Chromatogram
1.140
1.57%
1.253
0.33%
1.368
98.10%
Retention time (min)
/Users/sellmye...SS_Tz_Mor4.raw/ Injection 1 MS ES+ TIC
0.330
1.159
1.260
1.379
1.886
2.055
2.208
2.301
2.470
2.647
2.783
2.944
Retention time (min)
/Users/sellmye...SS_Tz_Mor4.raw/ Injection 1 MS ES+ MS + spectrum 1.37
120.089
7.19%
157.147
4.28%
201.260
1.20%
270.286
1.42%
288.325
24.21%
289.333
3.96%
343.446
1.31%
357.666
63.85%
358.386
100.00%
359.394
13.89%
379.339
2.08%
481.540
0.59%
m/z (Da)

SS_Tz_NNDM.12.fid

### $^{1}H$ NMR, 600 MHz, DMSO-$d_6$

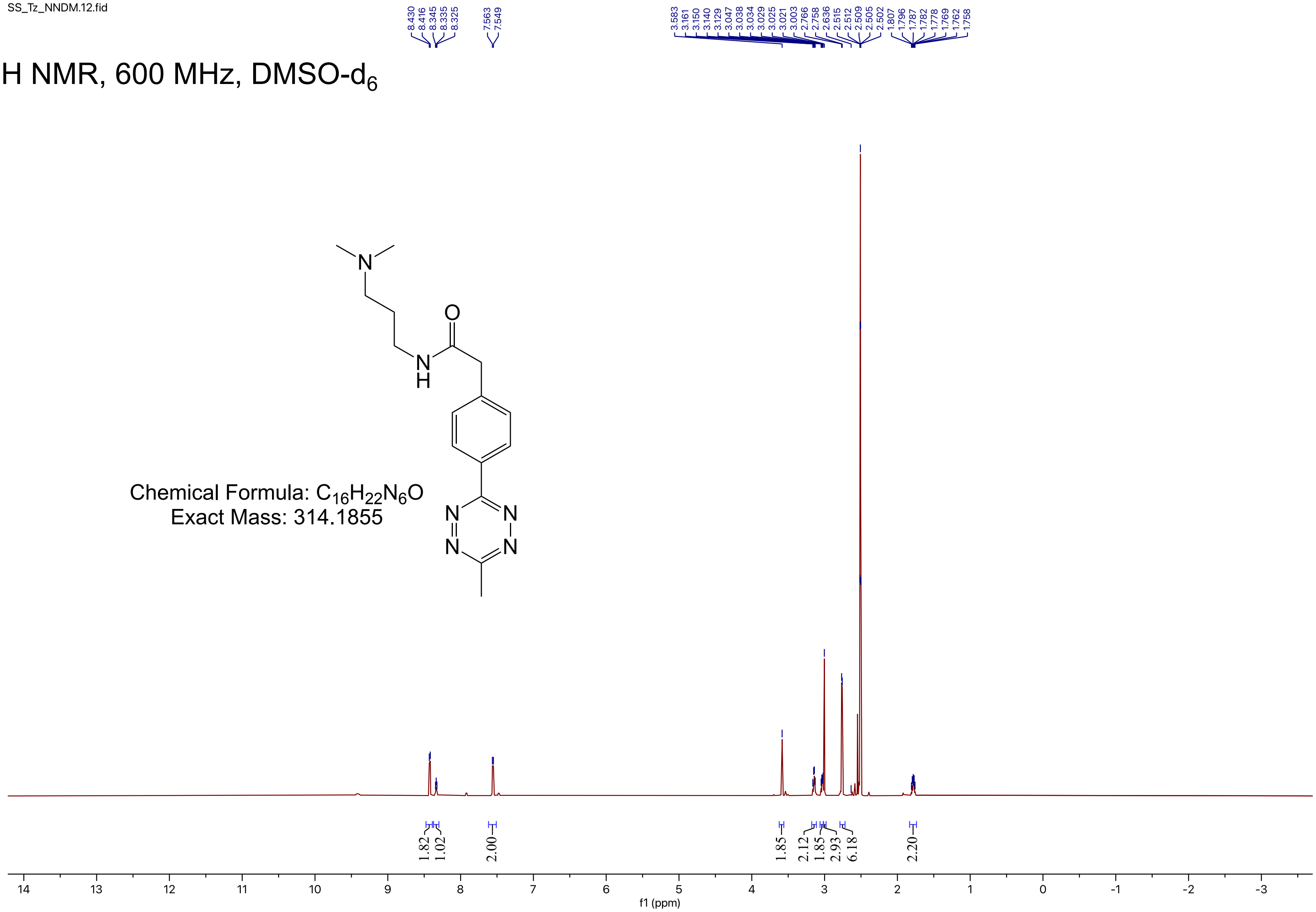


Chemical Formula: $C_{16}H_{22}N_6O$
Exact Mass: 314.1855

SS_Tz_NNDM_13C.1.fid

### $^{13}C$ NMR, 100 MHz, DMSO-$d_6$

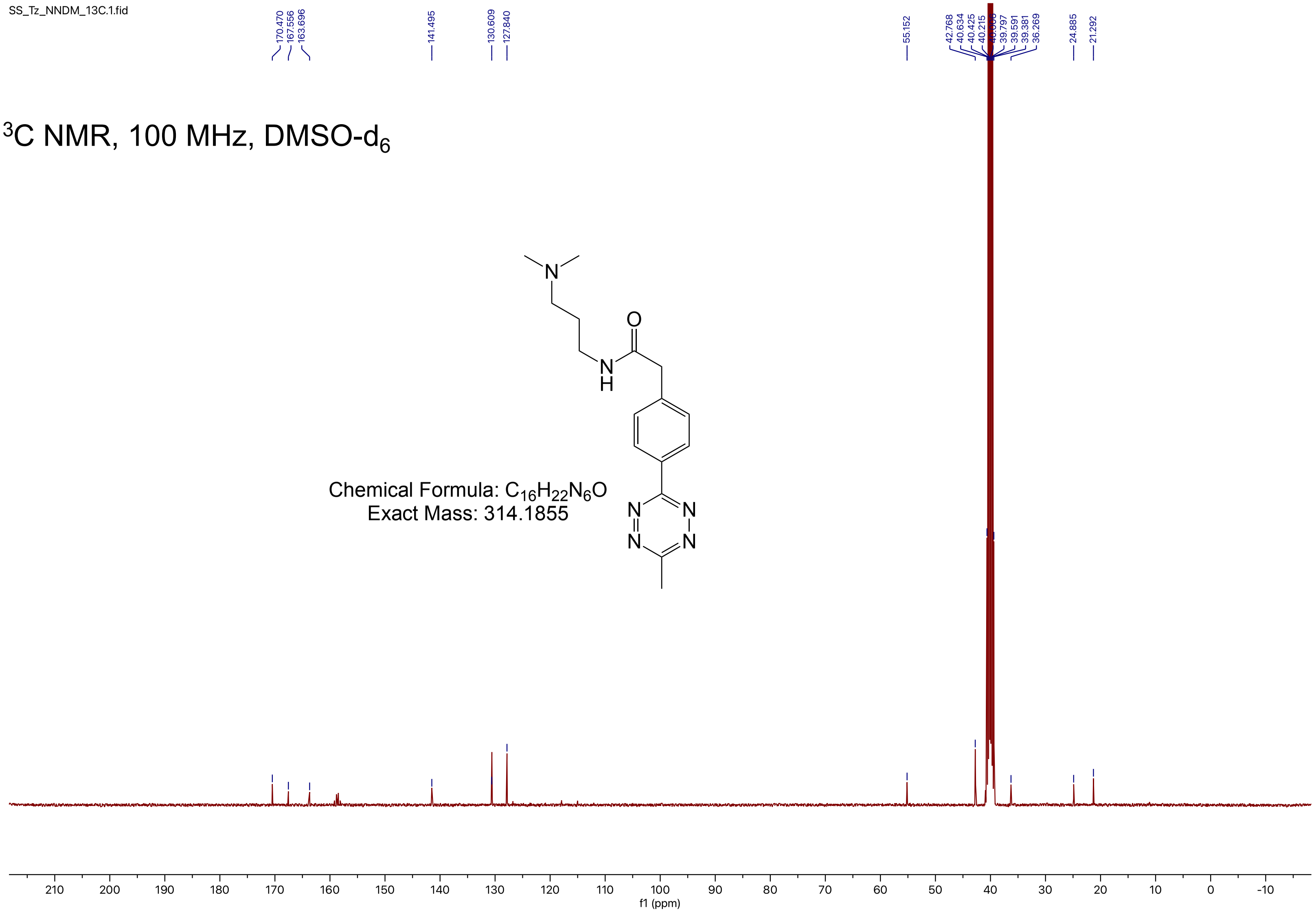


Chemical Formula: $C_{16}H_{22}N_6O$
Exact Mass: 314.1855

D:\Data\Sellmyer\SS_NMDM 11/02/23 16:08:08

### HRMS

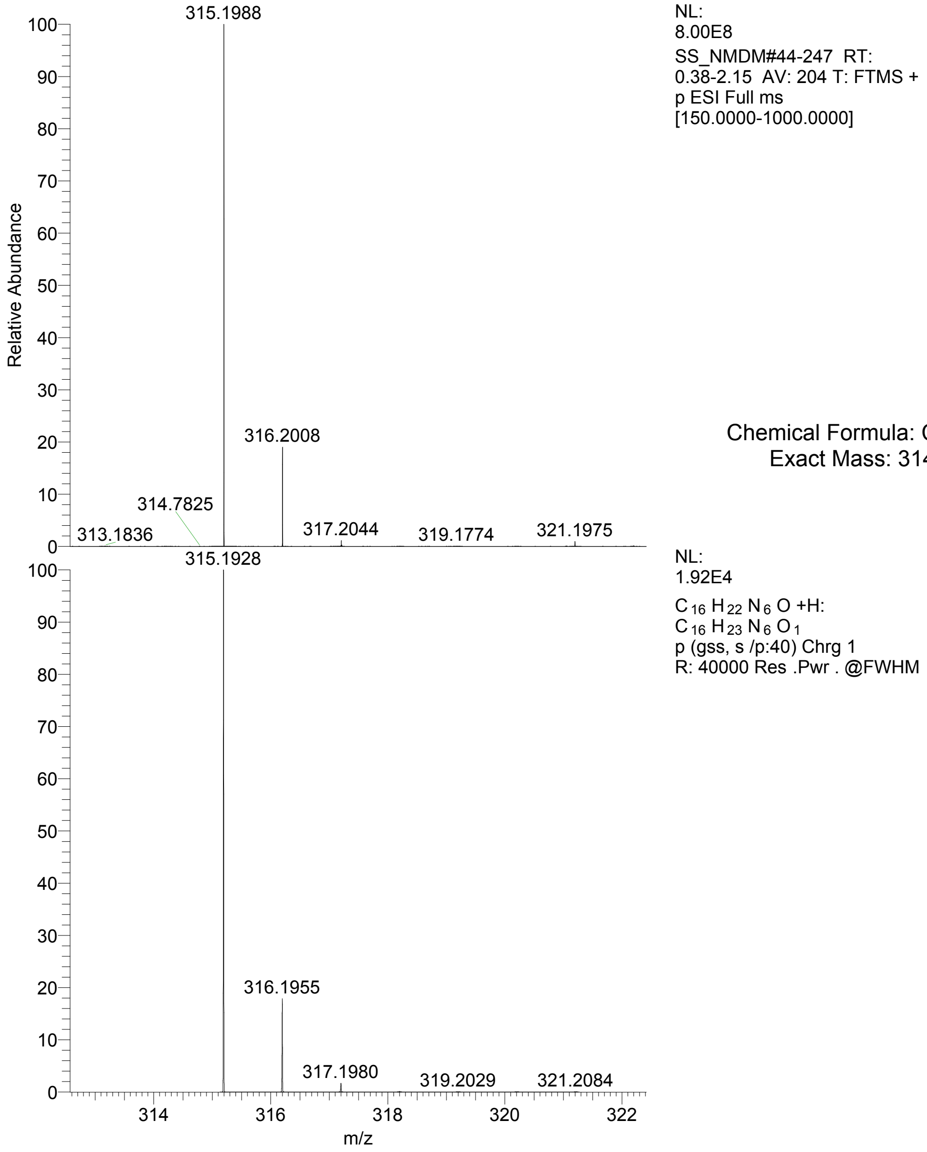


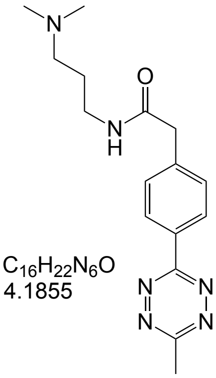

Chemical Formula: $C_{16}H_{22}N_6O$
Exact Mass: 314.1855

### LCMS

Chemical Formula: $C_{16}H_{22}N_6O$
Exact Mass: 314.1855

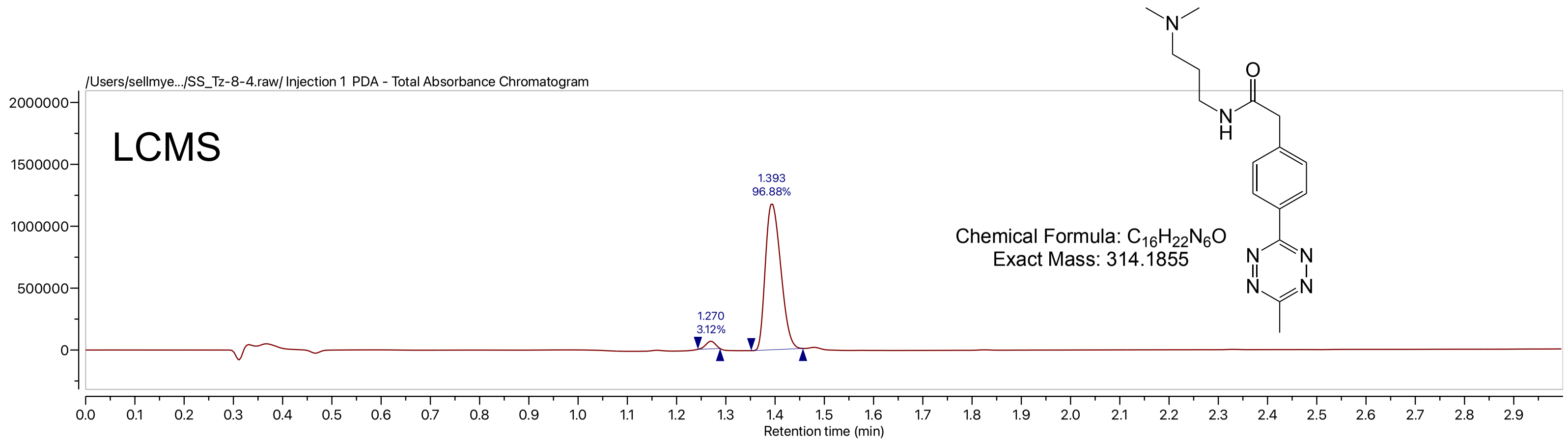


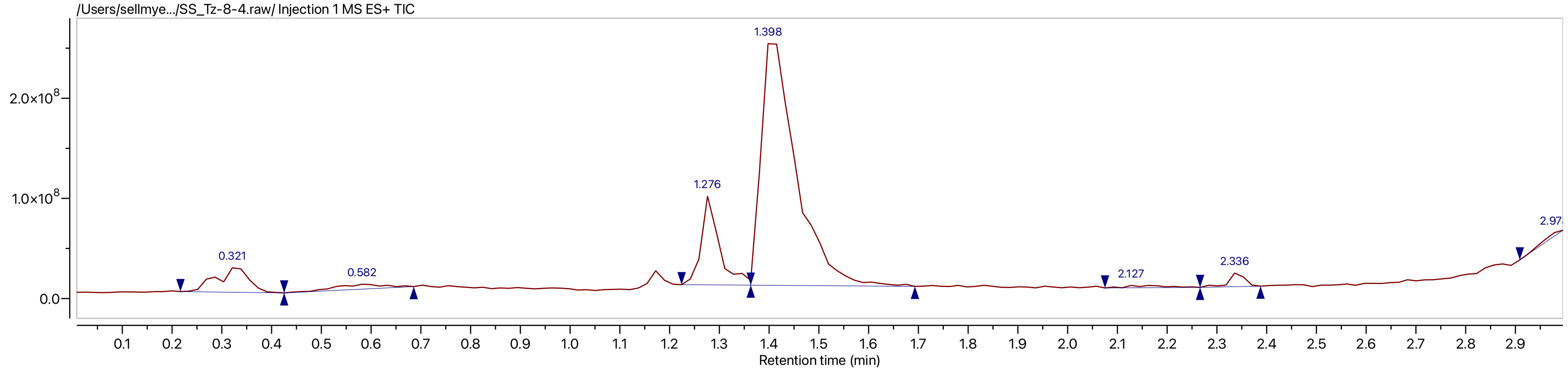


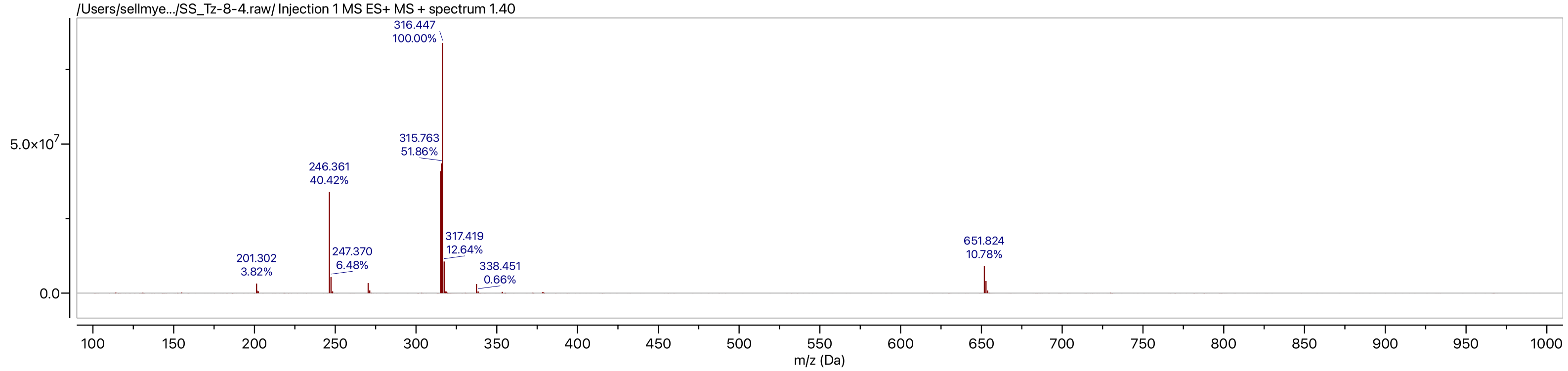

SS_Tz_PEG.1.fid

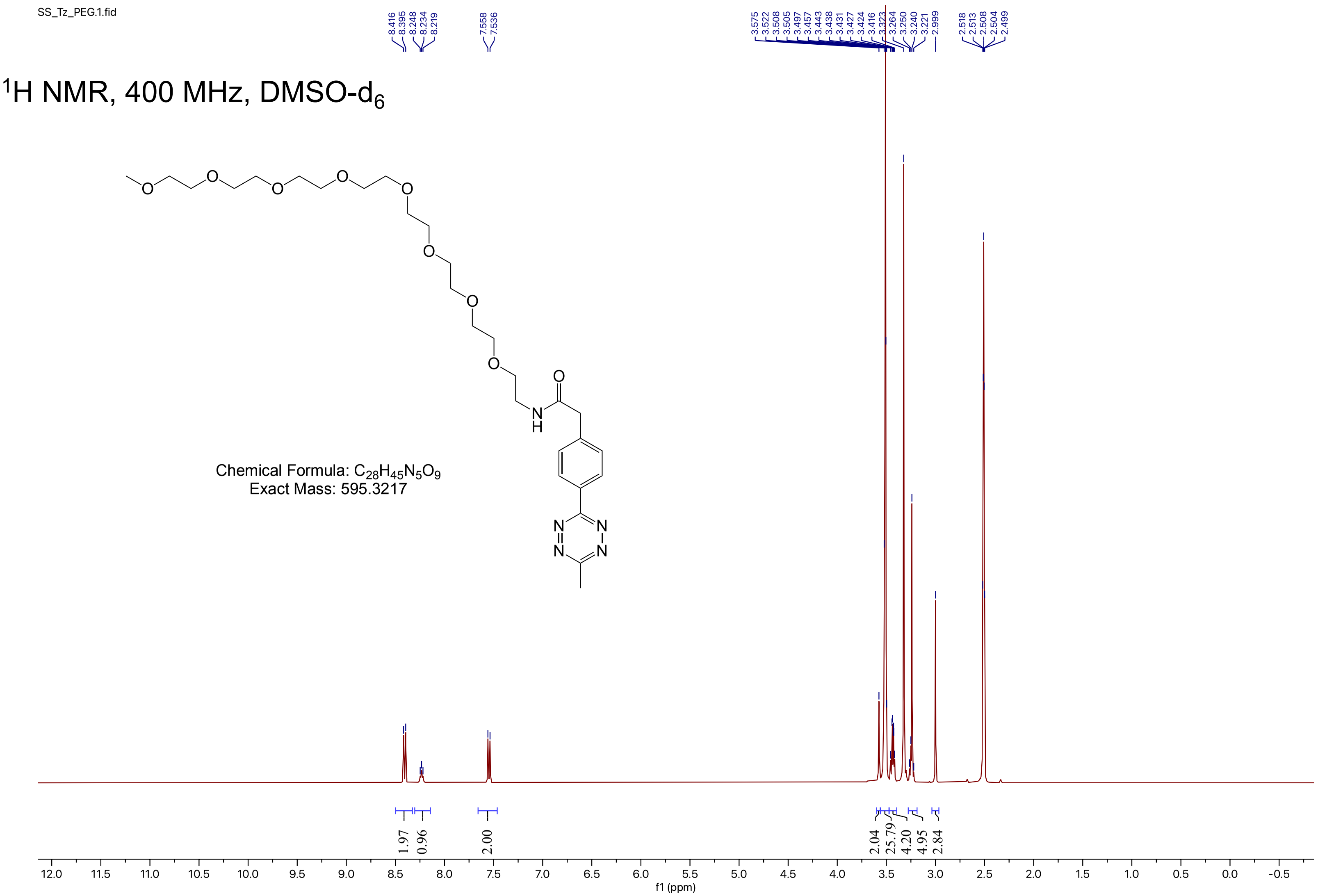

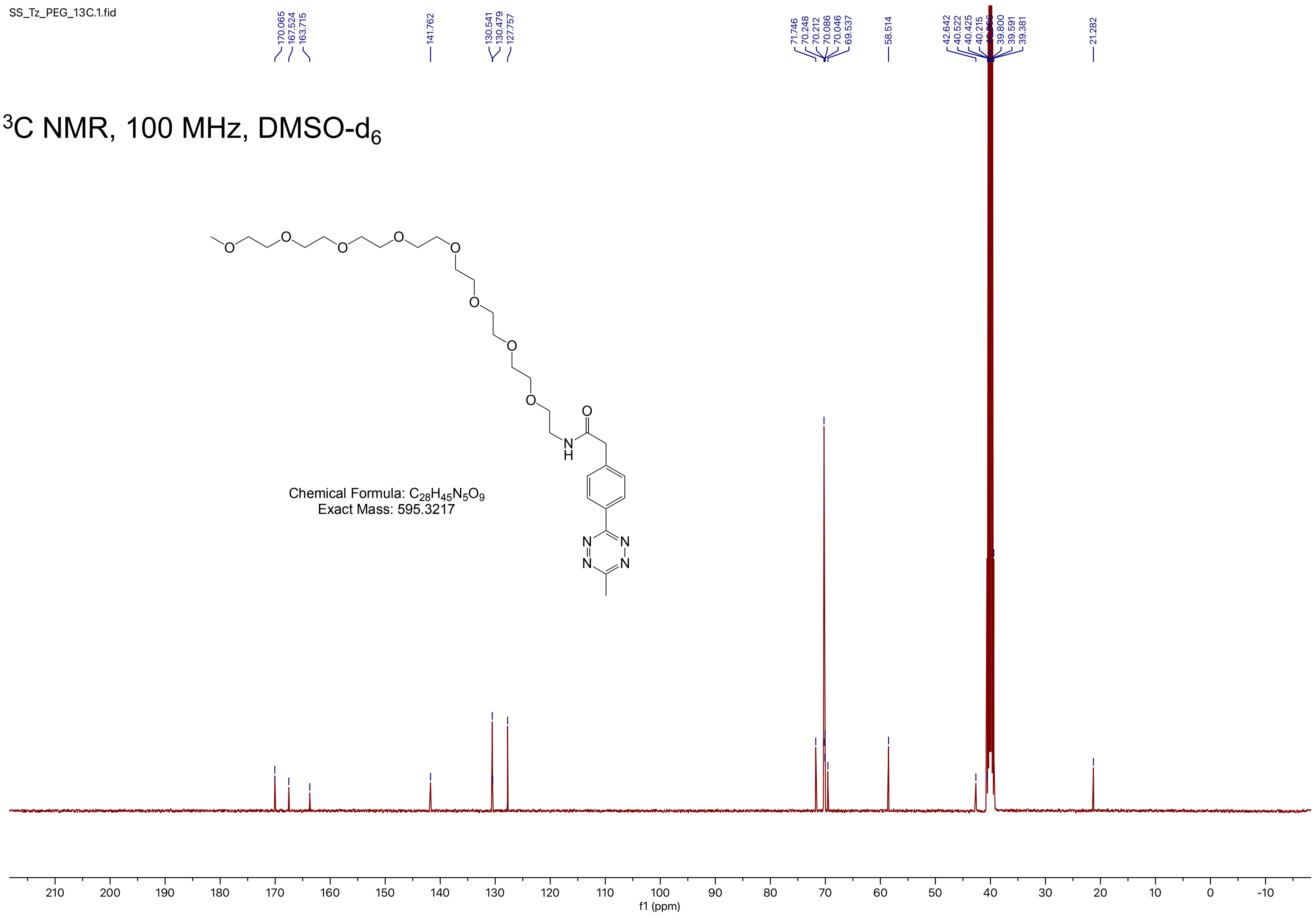
SS_Tz_PEG_13C.1.fid
170.065
167.524
163.715
141.762
130.541
130.479
127.757
71.746
70.248
70.212
70.086
70.046
69.537
58.514
42.642
40.522
40.425
40.215
[illegible]
39.800
39.591
39.381
21.282
$^{13}C$ NMR, 100 MHz, DMSO-$d_6$
Chemical Formula: $C_{28}H_{45}N_5O_9$
Exact Mass: 595.3217
210
200
190
180
170
160
150
140
130
120
110
100
90
80
70
60
50
40
30
20
10
0
-10
f1 (ppm)

D:\Data\Sellmyer\SS_PEh 11/02/23 15:31:47

### HRMS

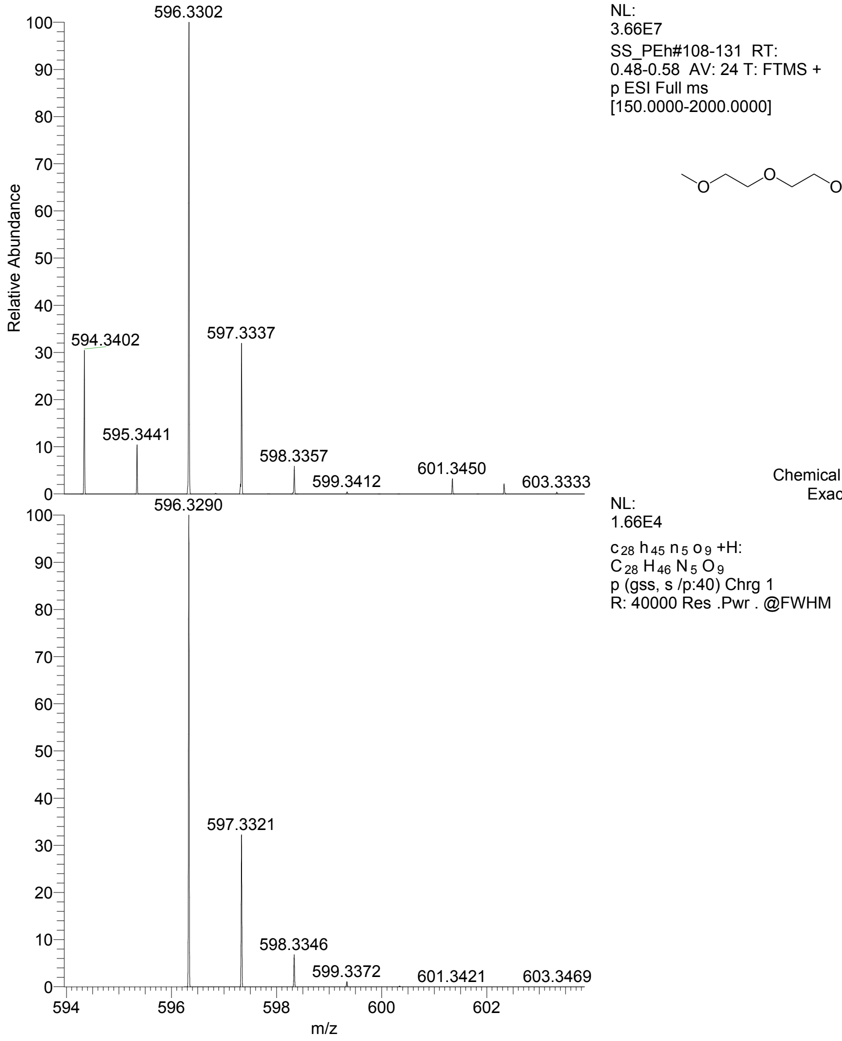


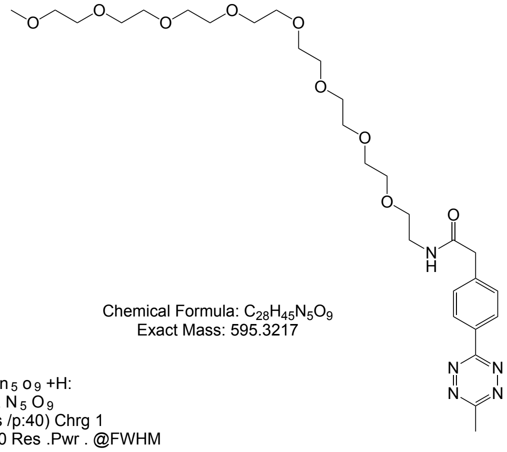

Chemical Formula: $C_{28}H_{45}N_5O_9$
Exact Mass: 595.3217

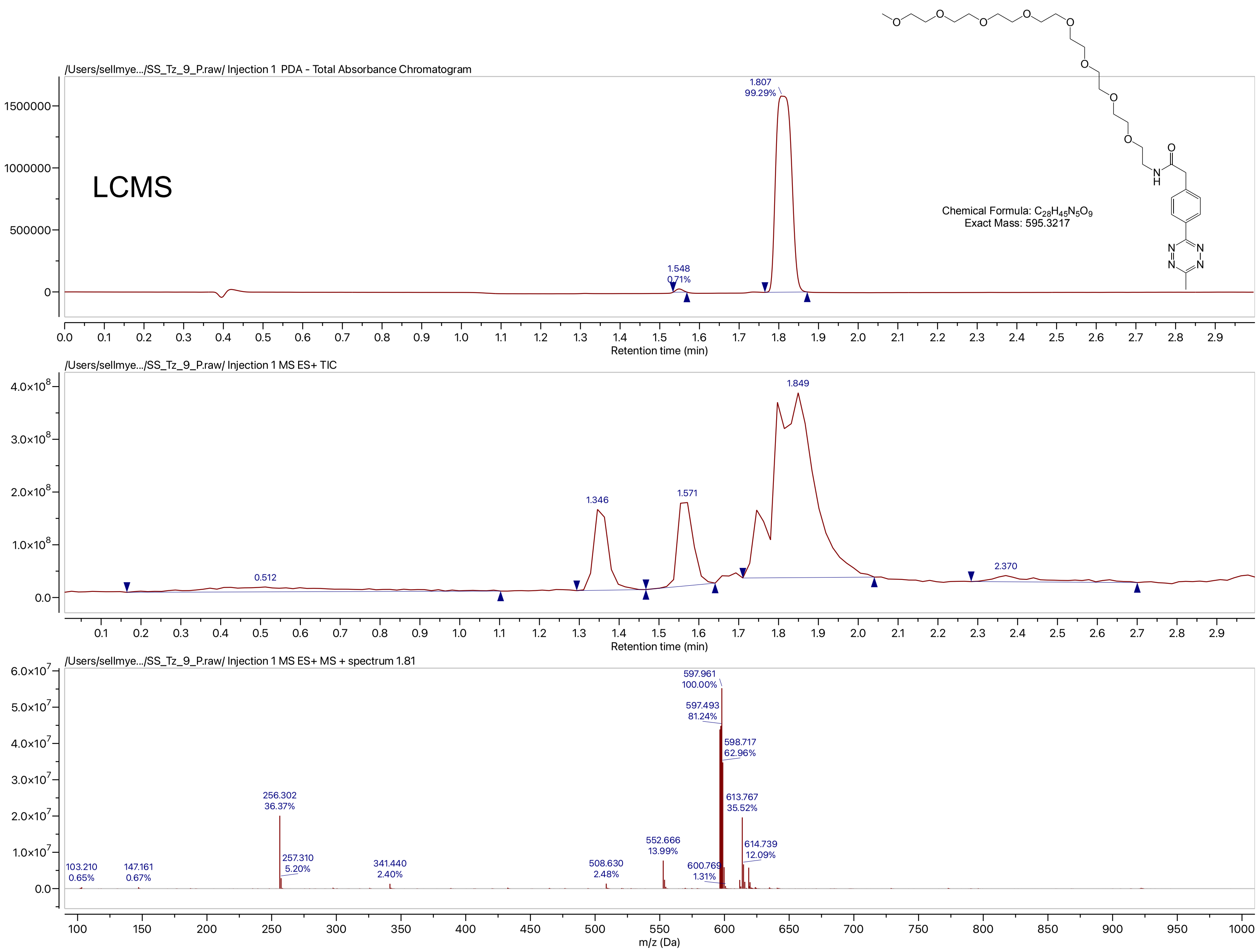
LCMS
/Users/sellmye.../SS_Tz_9_P.raw/ Injection 1 PDA - Total Absorbance Chromatogram
1.807
99.29%
1.548
0.71%
Retention time (min)
Chemical Formula: C28H45N5O9
Exact Mass: 595.3217
/Users/sellmye.../SS_Tz_9_P.raw/ Injection 1 MS ES+ TIC
0.512
1.346
1.571
1.849
2.370
Retention time (min)
/Users/sellmye.../SS_Tz_9_P.raw/ Injection 1 MS ES+ MS + spectrum 1.81
103.210 0.65%
147.161 0.67%
256.302 36.37%
257.310 5.20%
341.440 2.40%
508.630 2.48%
552.666 13.99%
597.961 100.00%
597.493 81.24%
598.717 62.96%
600.769 1.31%
613.767 35.52%
614.739 12.09%
m/z (Da)
